# Integrated multi-omic analysis of pediatric metastatic osteosarcoma reveals endothelial cell plasticity and lineage infidelity

**DOI:** 10.64898/2026.08.11.744222

**Authors:** Julian Burks, Ying Wu, Krithika Bhuvaneshwar, Neeraja Syed, Dahee Jung, Carly M. Sayers, Demond O. Williams, Swapna Vidhur Daulatabad, Therry Malone, Julie Galindo, Monica Mendez, Jennifer Cotter, Jovana Pavisic, Yoh-Suke Mukouyama, Jack F. Shern, Rosandra N. Kaplan, Troy A. McEachron

**Author notes:** These authors contributed equally to this work (Julian Burks, Ying Wu). **Corresponding Author:** Troy A. McEachron Pediatric Oncology Branch, Center for Cancer Research, National Cancer Institute, National Institutes of Health 10 Center Drive, Bethesda, MD 20892.

## Abstract

While recent research has increasingly focused on the role of fibroblasts and macrophages in osteosarcoma, the tumor vasculature remains poorly understood, particularly in metastatic disease. To address this gap, we performed single-nuclei multi-ome (RNA+ATAC) sequencing on 24 human metastatic osteosarcoma specimens. We found that endothelial cells adopt a hybrid endothelial-mesenchymal state resembling endothelial-to-mesenchymal transition (EndMT) and that a subset of diploid endothelial cells expresses osteoblastic transcriptional profiles and gene regulatory networks (GRN). Joint copy-number analysis further identified osteosarcoma cells with endothelial transcriptional programs and GRNs, consistent with vascular mimicry. *In vitro* assays and syngeneic lineage-tracing experiments validated that tumor educated endothelial cells acquire osteoblast-like features. Together, these findings reveal substantial plasticity among endothelial and osteosarcoma cells in human and murine metastatic osteosarcoma, provide new insight into the how the metastatic microenvironment shapes the tumor vasculature, and challenge current models of osteosarcoma biology.

## INTRODUCTION

Osteosarcoma is the most common malignant bone tumor in children, adolescents, and young adults (1). Metastatic disease is the strongest adverse prognostic factor as patients who present with, or develop metastases, have a median overall survival of less than two years and a 5-year overall survival probability of approximately 30% (2, 3). Despite decades of clinical trials, outcomes and treatment options for metastatic osteosarcoma have changed little over the past ∼40 years (1, 2, 4, 5). This plateau in progress underscores the need for deeper functional and biological insight into osteosarcoma metastases to enable new, mechanism-based treatment strategies.

A pathological diagnosis of osteosarcoma requires the presence of malignant osteoid and/or chondroid matrix produced by tumor cells. In the context of osteosarcoma pulmonary metastases, the ability of these tumors to deposit malignant osteoid/chondroid matrix requires the generation of a supportive osteoinductive microenvironment at the secondary disease site. Recent descriptive and functional studies have begun to map the molecular and cellular ecosystems of primary and metastatic osteosarcoma, with particular emphasis on fibroblasts and macrophages (6–18). In contrast, the vascular compartment of metastatic osteosarcoma, and how an osteogenic metastatic niche shapes tissue-resident vascular cells, remains largely unexplored.

To address this gap, we applied 10x Genomics single-nuclei multiome (snMulti-ome) profiling to characterize the vasculature across a cohort of 24 pediatric metastatic osteosarcoma specimens. Our data shows that endothelial cells consistently adopt a hybrid endothelial–mesenchymal state, and that a subset of these cells transition towards an osteoblast-like state. We also identify a small population of osteosarcoma cells that activate endothelial transcriptional and regulatory programs, consistent with vascular mimicry. Functional studies demonstrate that endothelial cells acquire osteoblastic phenotypic traits in response to tumor-derived cues both *in vitro* and *in vivo*. Together, these findings reveal extensive plasticity in both endothelial cells and osteosarcoma cells within metastatic disease.

## RESULTS

### Identification and subsetting of endothelial cells in the snMulti-ome dataset

The goal of this study was to use multi-omic profiling to better understand endothelial cell biology in metastatic osteosarcoma. We performed snMulti-ome sequencing on 24 frozen specimens from pediatric patients with pathologically confirmed metastatic osteosarcoma (**Supplemental Table 1**). This assay captures both RNA-seq and ATAC-seq from individual nuclei, enabling independent assessment of transcriptional programs and chromatin accessibility, as well as integrated analysis to better define cell state transitions. Cell-type annotation of the integrated dataset identified nuclei from endothelial cells, erythroid precursors, fibroblasts, lung epithelial cells, myeloid cells, cells of neuronal lineage, osteoclasts, osteosarcoma cells, smooth muscle cells, T cells, and an unassigned population that could not be definitively classified (**Figure 1A–B**). Nuclei from osteosarcoma cells, myeloid cells, endothelial cells, and fibroblasts were the most abundant populations (**Figure 1C**). Except for two specimens showing selective enrichment of nuclei from erythroid precursors or neuronal-lineage cells, these populations were otherwise present at low abundance across specimens (**Supplemental Figure 1**). Nuclei from lung epithelial cells were detected at low levels in most samples, with four specimens showing a substantial epithelial component (**Supplemental Figure 1**). Consistent with prior observations of lymphocyte exclusion in metastatic osteosarcoma (7, 9, 14), T cells were rare, comprising only 0.9% of nuclei (**Figure 1A, C; Supplemental Figure 1**).

**Figure 1:**
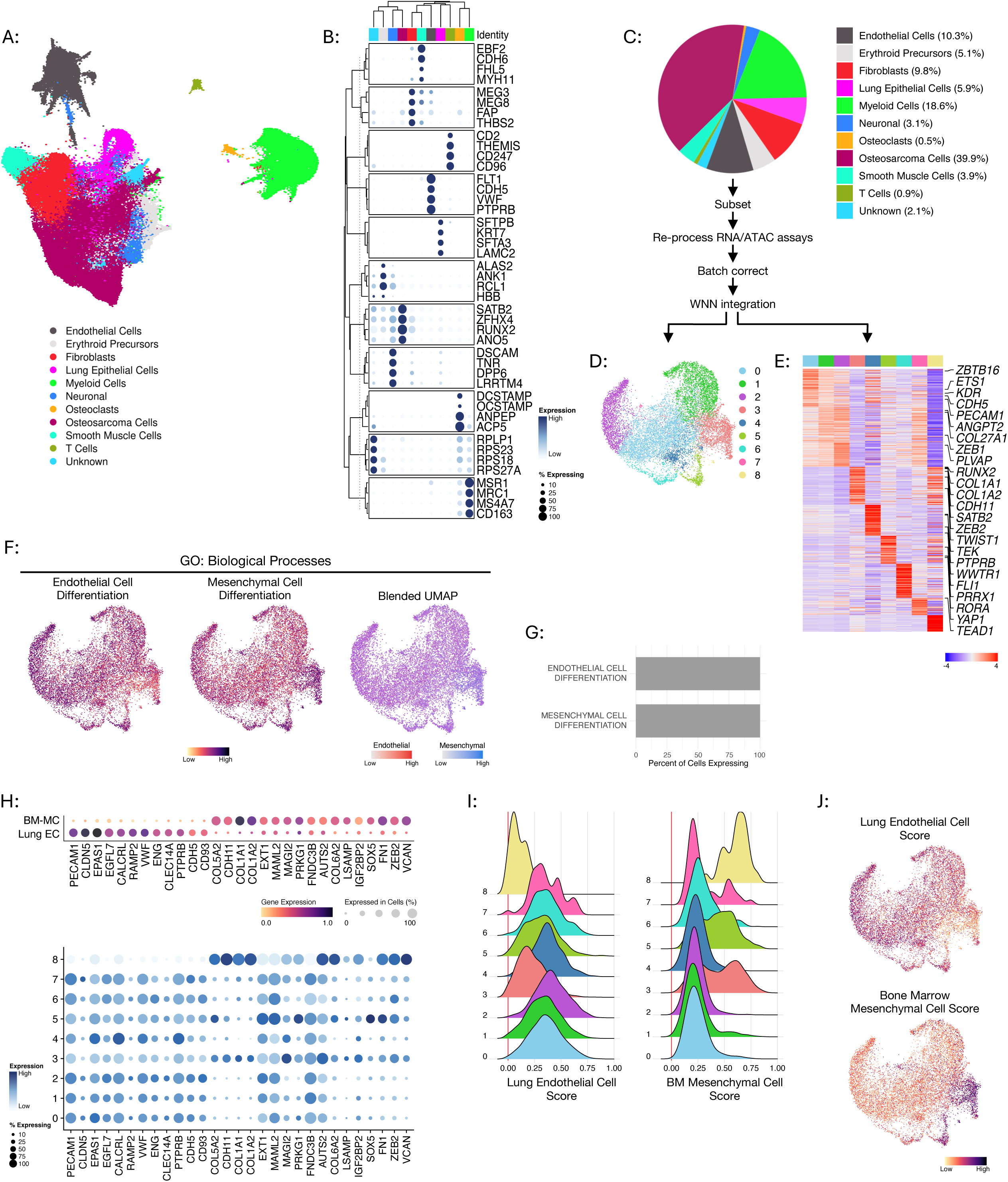
Tumor associated endothelial cells express mesenchymal lineage genes and metagene signatures indicative of a hybrid cell state. (A-C) Characterization of the snMulti-ome dataset. (A) WNN UMAP of the annotated dataset. (B) Clustered dot plot of marker genes for each cellular annotation according to the colors shown in panel (A) to the left. (C) Pie chart showing the relative proportions of each cellular annotation in the dataset. The workflow for subsetting and reprocessing the endothelial cells from the dataset is shown below the pie chart. (D) UMAP of the subsetted and re-processed endothelial cell nuclei. (E) Heatmap showing the cluster-specific highly variable genes with genes of interest annotated to the right. Color bar represents z-scaled values (blue = -2, red = +2). The color palettes in between panels (D) and (E) are consistent and denote the same clusters in each panel. (F) Results of gene set enrichment analysis projected on to UMAP plots. The endothelial cell differentiation signature (left) and the mesenchymal cell differentiation signature (middle) were scored using the Gene Ontology Biological Processes collection in MSigDb. Right: Blended UMAP overlaying the endothelial cell differentiation (red) and mesenchymal cell differentiation (blue) enrichment scores. Gradations of purple indicates presence of both signatures in the same nucleus relative to each score. (G) Quantification of the percentage of nuclei expressing the endothelial cell differentiation (top) and mesenchymal cell differentiation (bottom) signatures from the mSigDb Gene Ontology Biological Processes collection. (H) Dot plots of bone marrow mesenchymal cell (BM-MC) and lung endothelial cell (Lung-EC) marker genes in the Tabula Sapiens atlas (top) and this dataset (bottom). (I) Ridge plots showing the cluster-specific distribution of the Lung-EC (left) and BM-MC (right) gene signatures created from the Tabula Sapiens dataset. Each cluster is colored according to the UMAP shown in (D). (J) Projection of the Lung-EC (top) and BM-MC (bottom) metagene signature scores onto UMAP plots of the subsetted endothelial cell nuclei.

Weighted nearest neighbors (WNN) analysis is a computational method that integrates multiple single cell/nuclei profiling modalities (19). We used WNN analysis to integrate the gene expression and chromatin occupancy profiles to analyze nuclei based on cellular state. Clustering based on WNN integration better separated the clusters in 2-dimensional space and yielded a more distinct endothelial compartment than clustering using RNA alone (**Supplemental Figure 1A**). Canonical endothelial markers, including *CDH5, VWF, PTPRB,* and *CLDN5*, were almost exclusively expressed within the endothelial population (**Supplemental Figure 2**). Additional genes such as *ZEB1, TCF4*, and *BMPR2* were enriched in endothelial cell nuclei but also detectable at lower levels in other populations (**Supplemental Figure 2**). After subsetting the endothelial cluster, reprocessing the data, batch correcting, performing WNN integration, and re-clustering, we identified nine endothelial clusters (**Figure 1D**). Subsequent inspection revealed a well-integrated dataset free of overt batch effects with nuclei from the larger clusters present in each of the specimens and nuclei from the smaller clusters present in select specimens (**Supplemental Figure 3**). The subsetted endothelial cluster was subsequently analyzed as a separate dataset from the total dataset and was the focus of this study.

### Tumor-associated endothelial cells exhibit mesenchymal transcriptional programs

Differential gene expression analysis was performed to define the transcriptional programs underlying each endothelial cluster. While endothelial lineage genes were robustly expressed, mesenchymal markers were also consistently detected (**Figure 1E**). To evaluate this more systematically, gene set enrichment analysis was performed using the Gene Ontology: Biological Processes gene set collection in MSigDB to quantify the enrichment of the “Endothelial Differentiation” and “Mesenchymal Differentiation” signatures in individual nuclei. We rederived these signatures so that genes present in both signatures were removed to ensure that only genes unique to either process were being evaluated. Notably, both signatures were enriched in 100% of nuclei (**Figure 1F–G**). Doublets and multiplets were removed during the data preprocessing stage to avoid confounding interpretations of the results. In addition, profiling nuclei rather than intact cells reduces the likelihood that endothelial–mesenchymal heterokaryons could drive the observed transcriptional overlap.

To further investigate this observation using an independent reference, we derived bone marrow mesenchymal cell (BM-MC) and lung endothelial cell (Lung EC) signatures from non-diseased single-cell RNA-seq datasets in the Tabula Sapiens database and examined their expression across the nine endothelial cell clusters in our dataset. While the gene set enrichment analysis performed above focused on genes involved in the process of differentiation towards a mesenchymal or endothelial cell fate (**Figures 1F-G**), the BM-MC and Lung EC markers derived from the Tabula Sapiens database are indicative of terminally differentiated cells. Lung EC marker genes were strongly expressed in clusters 0–7 and minimally expressed in cluster 8, whereas BM-MC marker genes were most prominent in cluster 8 and also present across the remaining clusters (**Figure 1H**). We then generated metagene scores for both signatures. This analysis confirmed that all nuclei expressed both programs with clusters 3, 5, and 8 showing the strongest BM-MC enrichment (**Figure 1I–J**). These clusters can also reflect a continuum of differentiation with cluster 8 indicative of a more mesenchymal state. Together, these findings indicate that endothelial cells within metastatic osteosarcoma lesions exist in a hybrid cell state in which endothelial and mesenchymal programs are simultaneously active, consistent with an endothelial-to-mesenchymal EndMT-like phenotype (20).

### The intratumoral microenvironment is enriched in EndMT drivers

EndMT is regulated by multiple signaling pathways, including BMP, TGFβ, and WNT, whereas FGF2 has been reported to antagonize this process (21, 22). Based on our observation that endothelial cells in metastatic osteosarcoma specimens consistently adopt a hybrid endothelial–mesenchymal cell state, we next asked whether the metastatic osteosarcoma microenvironment supports an EndMT-permissive signaling state. To accomplish this, we used CytoSig to infer signaling activity across a cohort of nine metastatic osteosarcoma specimens previously profiled by Visium spatial transcriptomics (7, 23). Comparison of inferred signaling activity between intratumoral regions (tumor core) and extratumoral regions (tumor periphery) revealed that BMP6 and TGFβ1 activity were prominent features of the intratumoral microenvironment, whereas FGF2 activity was enriched in the extratumoral compartment (**Figure 2A–B**). Moderate WNT signaling activity was observed in both the intratumoral and extratumoral microenvironments. Consistent with this spatial pattern, FGF2 signaling activity was significantly lower than BMP6, TGFβ1, and WNT3A activity within intratumoral regions (**Figure 2C**).

**Figure 2:**
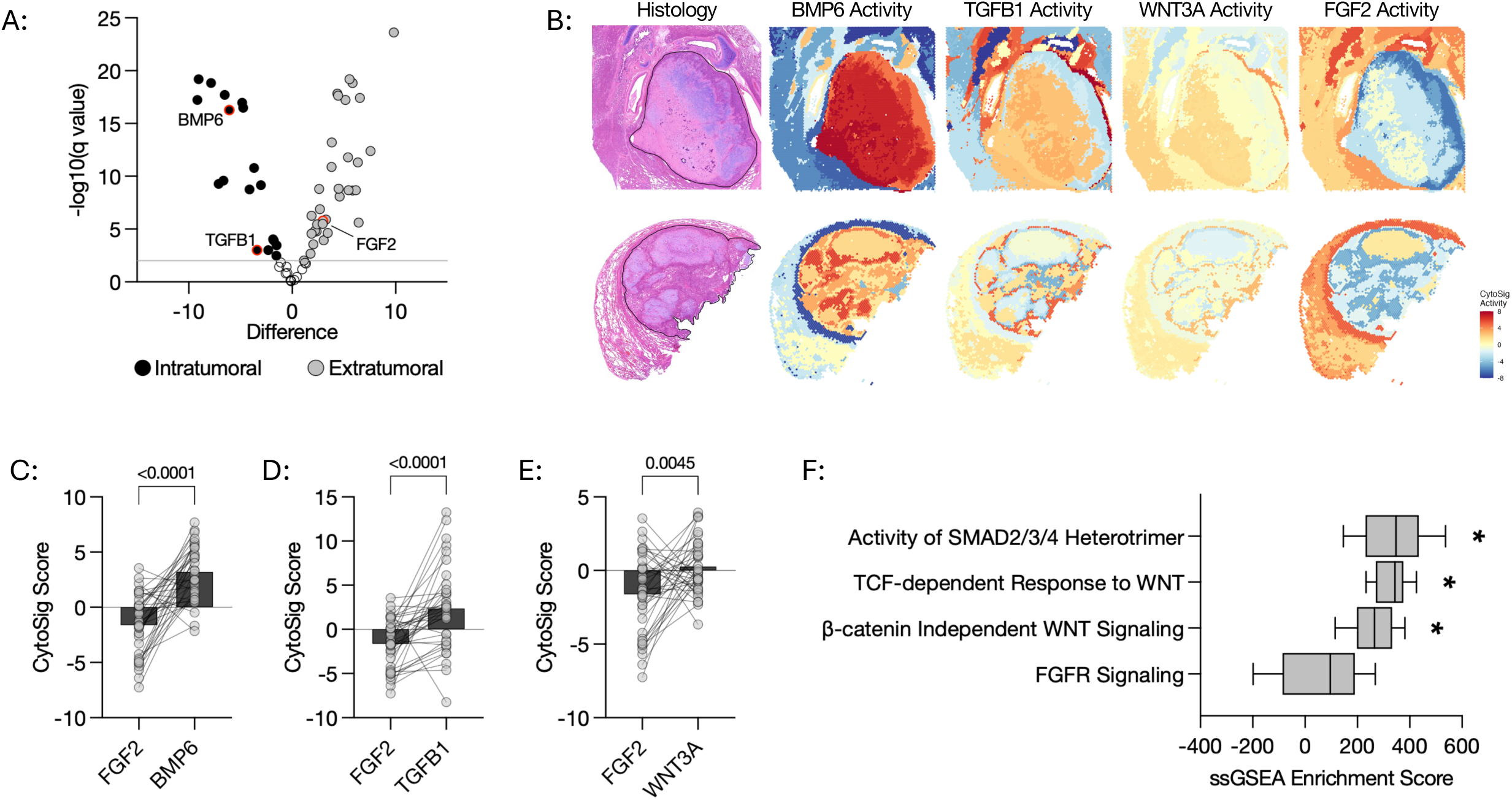
Spatial transcriptomic and bulk proteomic analysis of metastatic osteosarcoma specimens reveals an intratumoral signaling profile that favors EndMT. (A) Volcano plot showing the differentially enriched cytokine activity within the intratumoral microenvironment (gray circles) versus the extratumoral microenvironment (black circles) from a cohort of 9 metastatic osteosarcoma specimens profiled using the standard resolution Visium assay. Signaling molecules of interest are outlined in red and labeled accordingly. The horizontal gray line indicates p value threshold of 0.001. (B) Spatial mapping of the CytoSig activity scores in two representative metastatic osteosarcoma specimens. The malignant regions within each histological image are outlined in black and shown for reference. The intratumoral microenvironment corresponds to the region within the black outline and the extratumoral microenvironment is outside of the black outline. Color bar represents z-scaled values (blue = -8, red = +8). (C-E) Quantification of the differences in CytoSig activity scores between (C) FGF2 and BMP6, (D) FGF2 and TGFB1, and (E) FGF2 and WNT3A within the intratumoral microenvironment of osteosarcoma metastases. Each dot represents a spatial cluster in which the respective CytoSig activity was scored. Box plots show the aggregate mean of each score. Statistical significance determined using paired two-tailed t-tests where p < 0.05 is deemed significant. (F) Box plots showing the results of single sample gene set enrichment analysis of mass spectrometry data from a cohort of 10 metastatic osteosarcoma specimens. Reactome gene signatures are plotted on the Y-axis. Statistical significance determined using one-way ANOVA with a Dunnett’s multiple comparisons test comparing each gene signature to the FGFR Signaling signature. Asterisks indicate p values < 0.05.

To independently validate these findings, we performed single-sample gene set enrichment analysis using Reactome pathway signatures on mass spectrometry data generated from a subset of specimens used in this snMultiome profiling study. This orthogonal proteomic analysis demonstrated significantly reduced enrichment of FGFR signaling relative to SMAD and WNT signaling in these specimens (**Figure 2F**. These findings support the spatial profiling data demonstrating that WNT, BMP, and TGFB1 signaling are active within metastatic osteosarcoma specimens. Together, these data show that the intratumoral microenvironment of metastatic osteosarcoma is characterized by a signaling imbalance that favors EndMT and provides a molecular basis for the hybrid endothelial–mesenchymal cell state observed in tumor-associated endothelial cells.

### Identification of osteosarcoma vascular mimics

Vascular mimicry refers to tumor cells adopting endothelial-like transcriptional and phenotypic features (24). Given the increased BM-MC enrichment in clusters 3, 5, and 8 of the subsetted endothelial cell nuclei (**Figure 1H–J**), we wanted to determine if these clusters reflected endothelial cells further along an EndMT continuum or tumor cells exhibiting vascular mimicry. Osteosarcoma is characterized by extensive copy number alterations, providing a genomic feature that can help distinguish malignant from non-malignant diploid populations (25–28). Taking advantage of this osteosarcoma-intrinsic biomarker, we performed copy number inference independently using both the RNA assay (SCEVAN) and the ATAC assay (epiAneuFinder) and categorized each of the nuclei as tumor/aneuploid or normal/diploid (29, 30). Nuclei classified as diploid by both methods were labeled as “confident normal” and considered diploid endothelial cells, whereas nuclei classified as aneuploid were labeled as “confident tumor” and referred to as osteosarcoma vascular mimics (**Figure 3A**). Nuclei with discordant classifications were labeled “ambiguous”. Importantly, the “ambiguous” cells included those that were classified by SCEVAN yet were filtered out by epiAneuFinder as they failed to exceed the read count confidence threshold we established

**Figure 3:**
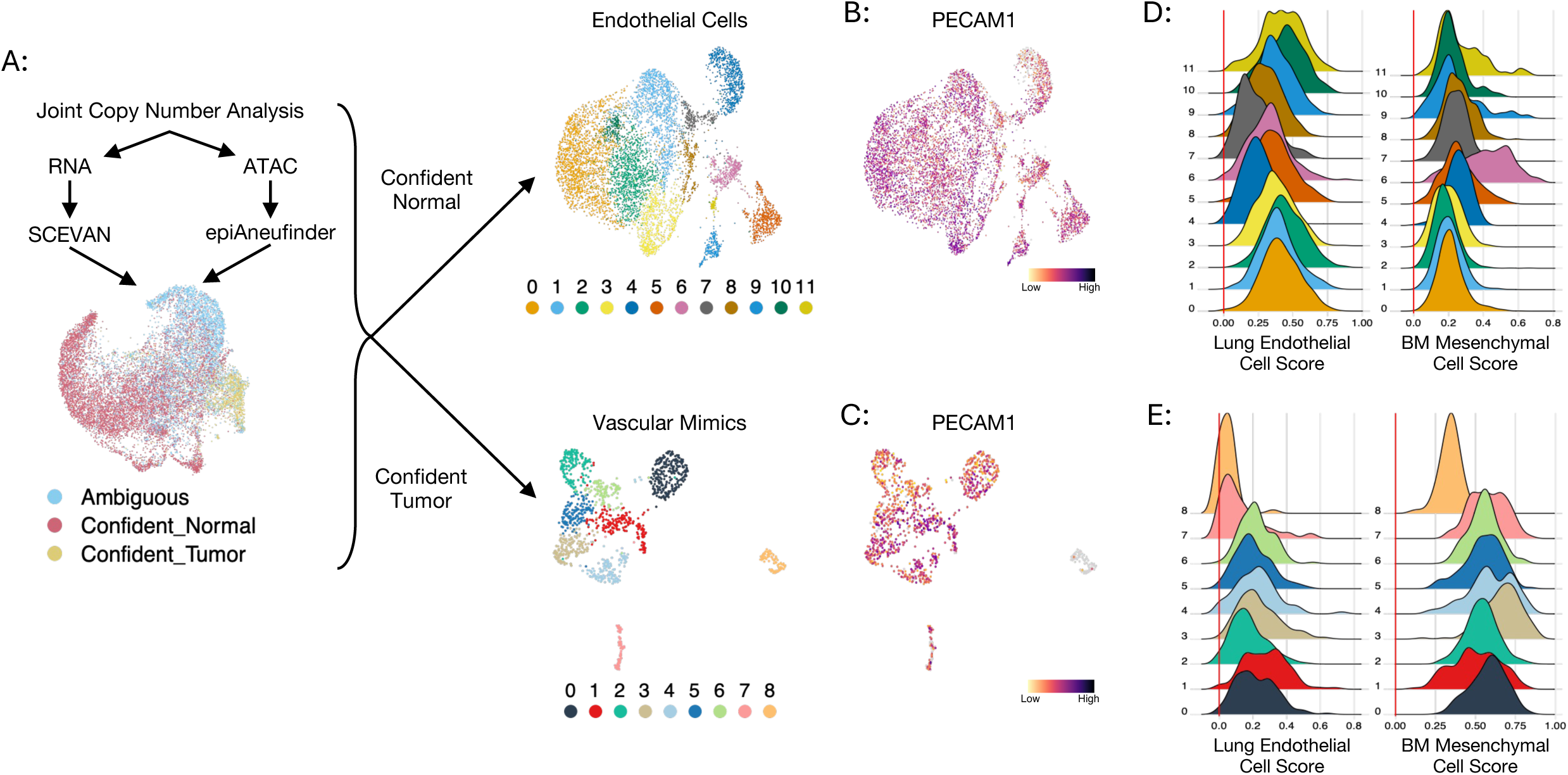
Joint copy number analysis identifies osteosarcoma vascular mimics. (A) Top: Schematic of the joint copy number calling analysis performed on both the gene expression (RNA) and ATAC (DNA) assays. Bottom: UMAP of the copy number analysis results for the subsetted endothelial cell nuclei using epiAneuFinder and SCEVAN. “Confident Normal” nuclei (diploid) are red, “Confident Tumor” nuclei (aneuploid) are yellow, and “Ambiguous” nuclei (low confidence nuclei and/or those below established thresholds) are blue. Right: UMAP plots of the subsetted and re-processed diploid endothelial cell nuclei (top UMAP) and osteosarcoma vascular mimic nuclei (bottom UMAP). The cluster assignments for each UMAP are indicated using a different color palette to denote separate data objects. (B-C) UMAP plots of *PECAM1* expression in (B) diploid endothelial cell and (C) osteosarcoma vascular mimic nuclei. (D-E) Ridge plots showing the cluster-specific distribution of the Lung-EC (left) and BM-MC (right) gene signatures in the (D) diploid endothelial cell nuclei and (E) osteosarcoma vascular mimics. Y-axis represents the number of nuclei within the given cluster assignment colored according to the UMAP plot shown in (A). Numbers along the y-axis represent the cluster identifier. Enrichment scores are plotted along the x-axis.

Next, we quantified the similarity within and between diploid endothelial cells, vascular mimics, and ambiguous endothelial populations to determine whether these represented distinct cellular subgroups. Although the nuclei showed high transcriptional correlation across groups, consistent with a shared endothelial identity, ATAC accessibility and WNN integration separated these populations more clearly (**Supplemental Figure 4**). Notably, diploid endothelial cells and vascular mimics exhibited limited similarity in both ATAC and WNN space, supporting their classification as distinct populations. In contrast, the ambiguous endothelial population showed greater similarity to both diploid endothelial cells and vascular mimics, suggesting that this group represents an admixed population (**Supplemental Figure 4**). These findings indicate that the endothelial populations differ more strongly at the chromatin regulatory level than at the steady-state RNA level, consistent with distinct regulatory networks, endothelial subtype identities, and/or activation states. This also highlights the utility of our joint copy-number and multiomic approach for resolving endothelial heterogeneity. To avoid confounding downstream analyses and inaccurate interpretation, nuclei from the ambiguous population were excluded from further analysis.

Osteosarcoma vascular mimics confidently accounted for 9.8% of the total endothelial subset (diploid endothelial cells, vascular mimics, and ambiguous populations) and were detected in 20 of the 24 specimens, with variable abundance across samples (**Supplemental Figure 5**). Nuclei from diploid endothelial cells and osteosarcoma vascular mimics were then separated, re-clustered, and analyzed independently (**Figure 3A**). The diploid endothelial cell nuclei resolved into 12 clusters, with cluster 4 composed almost entirely of nuclei from a metastatic liver lesion and expressed genes consistent with hepatic sinusoidal endothelium (**Figure 3A; Supplemental Figure 6**). Osteosarcoma vascular mimics partitioned into nine clusters. Cluster 8 was distinct in that it was well isolated on the UMAP and traced to a single specimen obtained from the superior sulcus, a highly innervated region at the lung apex (**Supplemental Figure 6**). Importantly, both diploid endothelial cells and osteosarcoma vascular mimics expressed the endothelial cel marker *PECAM1* and showed enrichment for both Lung EC and BM-MC metagene signatures across each of the clusters demonstrating that the hybrid endothelial-mesenchymal cell state is present in both groups (**Figure 3B–E**).

### GRN analysis and trajectory modeling of osteosarcoma vascular mimics

Vasculogenesis is the process of *de novo* vessel formation from precursor cells and is distinct from angiogenesis in which new vessels are formed from pre-existing vessels (31). Gene set enrichment analysis using the Gene Ontology: Biological Processes library revealed a uniformly high enrichment of the “Vasculogenesis” signature across the osteosarcoma vascular mimics (**Figure 4A**). To further characterize the nuclei from these cells and confirm the presence of endothelial cell regulatory programs, we analyzed transcription factor (TF) motif activity in the ATAC-seq data, focusing on TFs associated with endothelial identity. Cluster 8 was excluded from this analysis given that the entirety of this cluster is derived from a single specimen isolated from distinct and highly specialized tissue site (**Supplemental Figure 6**). Motif activity for canonical endothelial cell associated TF’s ETS1 and FLI1 were observed across most vascular mimic nuclei, with increased activity in clusters 1, 3, and 4 (**Figure 4B**). Similar motif activity patterns were observed for ETV2, a master regulator of vasculogenesis and endothelial identity (**Figure 4B**) (32). TF footprinting analysis further supported these findings, revealing increased chromatin accessibility flanking protected ETS1, FLI1, and ETV2 DNA binding motifs, consistent with TF occupancy at these loci (**Figure 4C**).

**Figure 4:**
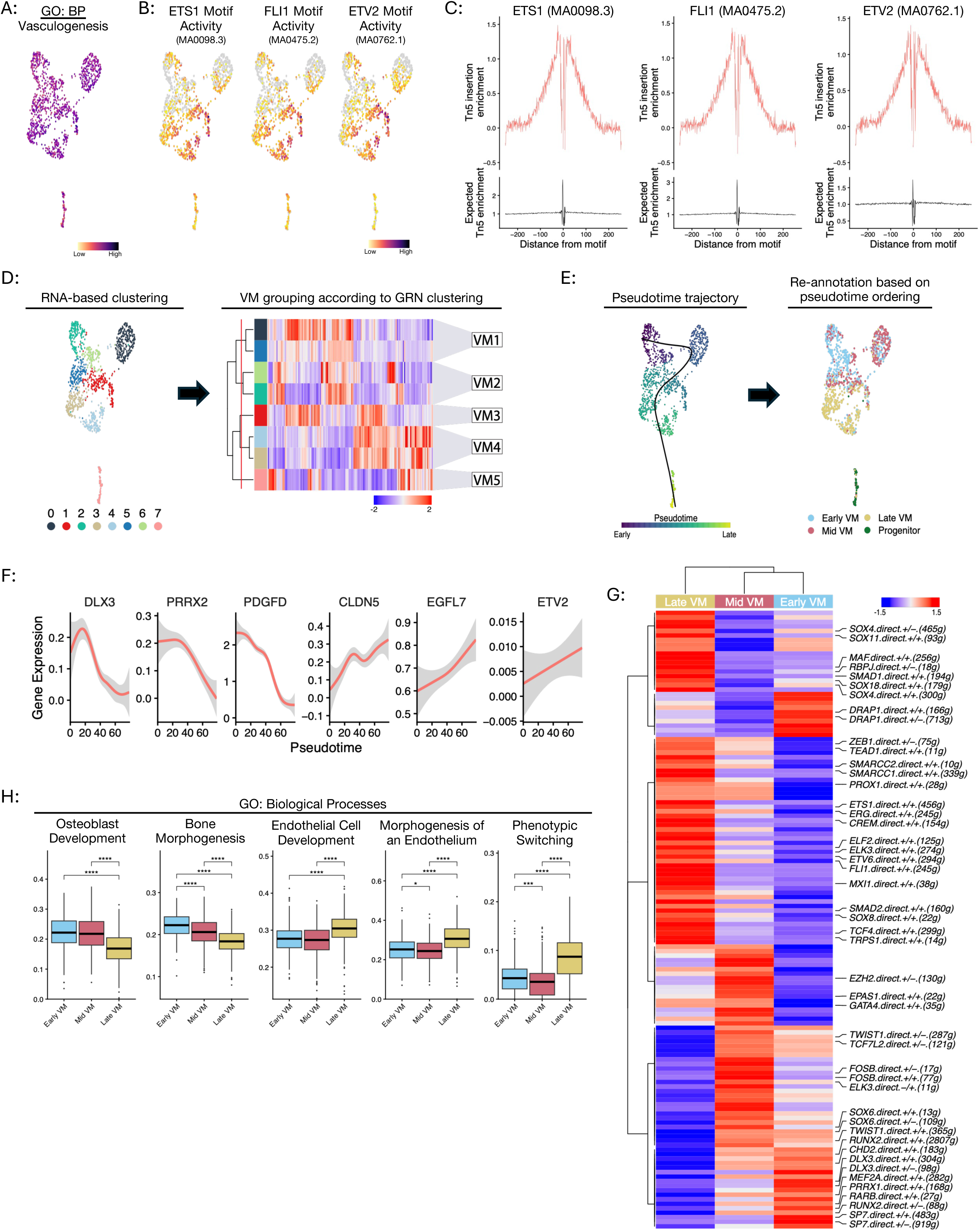
Molecular characterization of vascular mimicry in metastatic osteosarcoma. (A) UMAP results of gene enrichment analysis of the “Vasculogenesis” gene signature score from the Gene Ontology Biological Processes database. (B) UMAP plots of motif activity for TF’s associated with endothelial cell fate/identity and vascular mimicry. (C) TF footprinting profiles for ETS1 (left), FLI1 (middle), and ETV2 (right). (D) Heatmap showing the results of unsupervised hierarchical clustering (Spearman) of the GRNs identified in the osteosarcoma vascular mimics. The vertical red line indicates the dendrogram depth at which the clusters were reclassified into groups VM1-VM5. Color bar represents z-scaled AUC values (blue = -2, red = +2). Left: UMAP plot of the RNA-based clustering shown for reference. (E) Left: Pseudotime trajectory analysis projected on to the osteosarcoma vascular mimics UMAP. Color bar represents relative pseudotime values (blue = early, yellow = late). Right: The re-annotated UMAP plot showing cluster assignments according to the pseudotime ordering. (F) Trend curves showing changes in gene expression over pseudotime. (G) Heatmap showing the results of unsupervised hierarchical clustering (Spearman) of the GRNs identified in Early, Mid, and Late VM nuclei. Color bar represents z-scaled AUC values (blue = -2, red = +2). The signs following the GRN name reports the inferred activity: +/+ chromatin at locus is accessible and the target gene is expressed; +/- chromatin at locus is accessible and the target gene is not expressed; -/+ chromatin at locus is not accessible and the target gene is expressed; -/- chromatin at locus is not accessible and the target gene is not expressed. (H) Box plots of gene set enrichment scores in Early, Mid, and Late VM nuclei. Specific pathways are listed above each plot. Enrichment scores are plotted on the y-axis. Asterisks indicate statistically significant differences between the groups are determined by Wilcoxon signed rank test with Benjamini-Hochberg correction post-hoc test. P<0.001 considered statistically significant. Black bar indicates the mean value.

To better understand the regulatory programs associated with osteosarcoma vascular mimicry, GRNs were constructed linking TFs to their direct target genes. Unsupervised hierarchical clustering of the GRNs partitioned the nuclei into five groups, labeled VM1-VM5, representing different vascular mimicry cell states (**Figure 4D, Supplemental Figure 7**). Pseudotime analysis was then performed on the osteosarcoma vascular mimics to model the transdifferentiation trajectory (**Figure 4E**). The VM2 group was chosen as the starting point for this trajectory model as it consistently expressed the highest levels of osteoblast-related genes indicative of a more osteosarcoma-like cell state (**Supplemental Figure 8**). This analysis resulted in two pseudotime lineages (**Supplemental Figure 9**). Lineage 1 was comprised of VM1-VM4, whereas Lineage 2 was comprised of VM1-VM3, and VM5 (**Supplemental Figure 9**). Further analysis revealed an increased expression of the mesenchymal progenitor cell associated TF *PRRX1* in VM5 (**Supplemental Figure 9**). This data was interpreted as the nuclei in VM5 belonging to cells exhibiting a highly plastic and/or progenitor-like cell state. Assessment of developmental potency/stemness revealed that, while VM1-VM4 demonstrated similar degrees of stemness, VM5 scored significantly higher and was classified as multipotent, thus supporting VM5 as a progenitor-like population. Lastly, the nuclei were re-annotated according to their relative pseudotime ordering by dividing the pseudotime values into quartiles, thus partitioning the osteosarcoma vascular mimicry nuclei into the Early VM, Mid VM, Late VM and Progenitor groups (**Figure 4E**). The subsequent analyses focused on the Early VM, Mid VM, and Late VM groups of nuclei.

Analysis of gene expression changes in the osteosarcoma vascular mimics were consistent with a shift away from osteosarcoma/osteoblast identity over pseudotime, exemplified by decreased expression of osteoblast-associated genes *DLX3, PRRX2,* and *PDGFD* and increased expression of endothelial cell identity genes *CLDN5*, *EGFL7*, and *ETV2* (**Figure 4F**). Hierarchical clustering of GRNs showed distinct profiles between the Early, Mid, and Late VM groups with a selective enrichment of osteogenic GRNs in the Early VM group and increased expression of endothelial GRNs in the Late VM group (**Figure 4G**). Additionally, gene set enrichment analysis revealed decreased enrichment of osteogenic and bone-associated gene signatures and increases in endothelial gene signatures (**Figure 4H**). Interestingly, a significant enrichment in the “Phenotypic Switching” signature was observed in the Late VM group, implying that the process of osteosarcoma vascular mimicry is not unidirectional. Together, this integrated multi-omic analysis shows that a subset of metastatic osteosarcoma cells adopts endothelial-like transcriptional and gene regulatory programs consistent with vascular mimicry.

### Functional validation of osteosarcoma vascular mimicry in vitro and in vivo

Next we performed *in vitro* tube formation assays using four patient-derived osteosarcoma cell lines (33) to functionally determine if osteosarcoma cells can adopt endothelial phenotypic traits. Of the lines investigated, only OS052 and OS384 formed tube-like structures in wells coated with Matrigel basement membrane extract (**Figure 5A-B**). Interestingly, OS384 quickly formed unstable tubes that rapidly collapsed, whereas OS052 formed more stable tubes that persisted throughout the duration of the assay. OS186 and OS525 did not form tubes under these conditions (**Figure 5A-B**), underscoring heterogeneity in vascular mimicry potential across osteosarcoma models.

**Figure 5:**
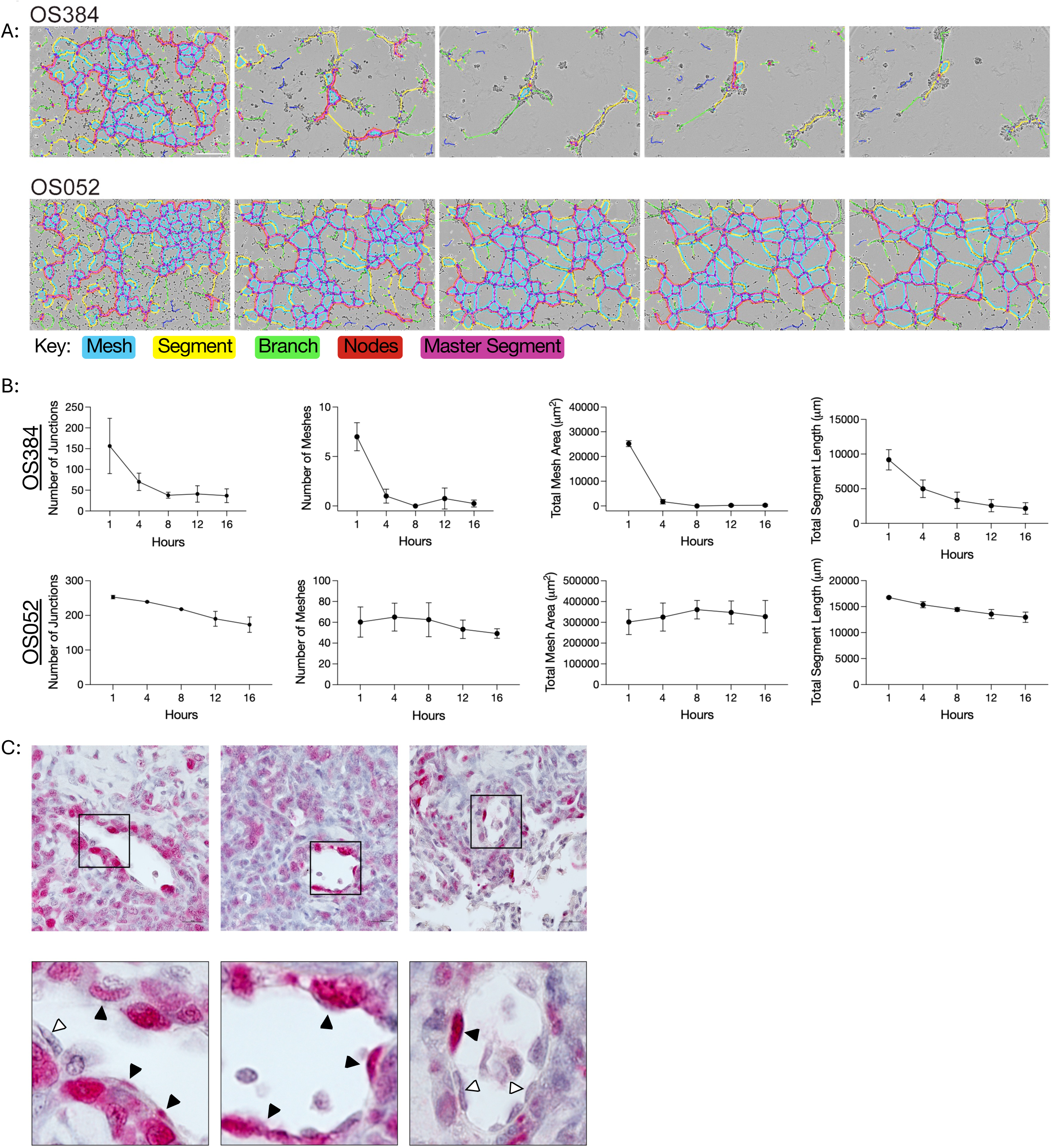
*In vitro* and *in vivo* validation of osteosarcoma vascular mimicry. (A) Representative brightfield images of Matrigel tube formation assays performed with OS384 and OS052 osteosarcoma cells, overlaid with network segmentation generated using the ImageJ Angiogenesis Analyzer plugin. Network components are color-coded as follows: segments (elements connecting two junctions), branches (elements connected to a single junction), junctions (nodes where three or more segments or branches intersect), meshes (fully enclosed network areas), master segments (segments connecting junctions), and total length (the cumulative length of all segments and branches). (B) Quantification of angiogenic network parameters generated by the Angiogenesis Analyzer at the indicated time points. Experiments were performed in duplicate. Data are presented as mean ± standard deviation. (C) Immunohistochemical analysis of vascular mimicry in spontaneous pulmonary metastatic lesions from the AXT syngeneic murine osteosarcoma model. GFP-expressing AXT osteosarcoma cells are shown in red. Insets highlight vascular structures exhibiting osteosarcoma vascular mimicry. Black arrowheads indicate luminal tumor cells, whereas white arrowheads indicate non-malignant cells lining the luminal surface. Scale bar, 20μm.

We further evaluated osteosarcoma vascular mimicry *in vivo* using the AXT immunocompetent syngeneic murine osteosarcoma model. This model is driven by *MYC* overexpression, *CDKN2A* loss, and mutant *TRP53* and is an appropriate phenocopy of human osteosarcoma as it forms spontaneous pulmonary metastases with malignant osteoid deposition (34). GFP-expressing tumor cells were identified in metastatic lung foci by immunohistochemistry using antibodies against GFP. Immunohistochemical analysis of the pulmonary metastatic lesions from the AXT model showed that GFP-positive tumor cells with elongated, spindle-shaped nuclei and cell bodies were integrated into the luminal surface of vascular structures (**Figure 5C**). Vascular mimicry events were rare, indicating that most vessels were comprised of endothelial cells and not vascular mimics. This is consistent with the snMulti-ome profiling data showing that osteosarcoma vascular mimics confidently comprised approximately 10% of the endothelial subset (**Supplemental Figure 5D**). Collectively, the computational and experimental results provide strong evidence that subsets of tumor cells in the metastatic osteosarcoma microenvironment undergo a phenotypic switch to mimic endothelial cells.

### Identification and characterization of osteogenic transdifferentiation of endothelial cells

Direct ossification, more commonly known as intramembranous ossification, is the physiological process in which mesenchymal cells differentiate directly into osteoblasts, bypassing the chondrocyte intermediate required for endochondral ossification (35). Given the enrichment of osteoinductive and EndMT-permissive signaling within the intratumoral osteosarcoma microenvironment, together with the consistent adoption of a hybrid endothelial–mesenchymal state by diploid endothelial cells (**Figures 1–3**), we hypothesized that these cells are primed to respond to osteosarcoma-derived osteoinductive cues and acquire osteogenic features. Because mesenchymal progenitor cells serve as precursors for intramembranous ossification, we next assessed whether these hybrid endothelial–mesenchymal cells showed evidence of osteoblastic priming or potential using the “Direct Ossification” gene signature from the MSigDB Gene Ontology collection as a surrogate transcriptional marker. To avoid confounding results from the osteosarcoma vascular mimics, we restricted these analyses to diploid endothelial cell nuclei and not the osteosarcoma vascular mimics.

Gene set enrichment analysis confirmed that all diploid endothelial nuclei were enriched in the “Mesenchymal Cell Differentiation” signature and that approximately 40% of nuclei were also enriched for the “Direct Ossification” signature (**Figure 6A-B**). Ridge plots were used to identify thresholds with which to stratify the diploid endothelial nuclei into Ossification Low, Ossification Med, and Ossification High groups based on the “Direct Ossification” gene set enrichment scores (**Figure 6C; Supplemental Figure 10**). Notably, most nuclei in cluster 4, composed almost entirely of nuclei from the metastatic liver lesion, was classified as Ossification Low (**Figure 6A, C; Supplemental Figure 6**).

**Figure 6:**
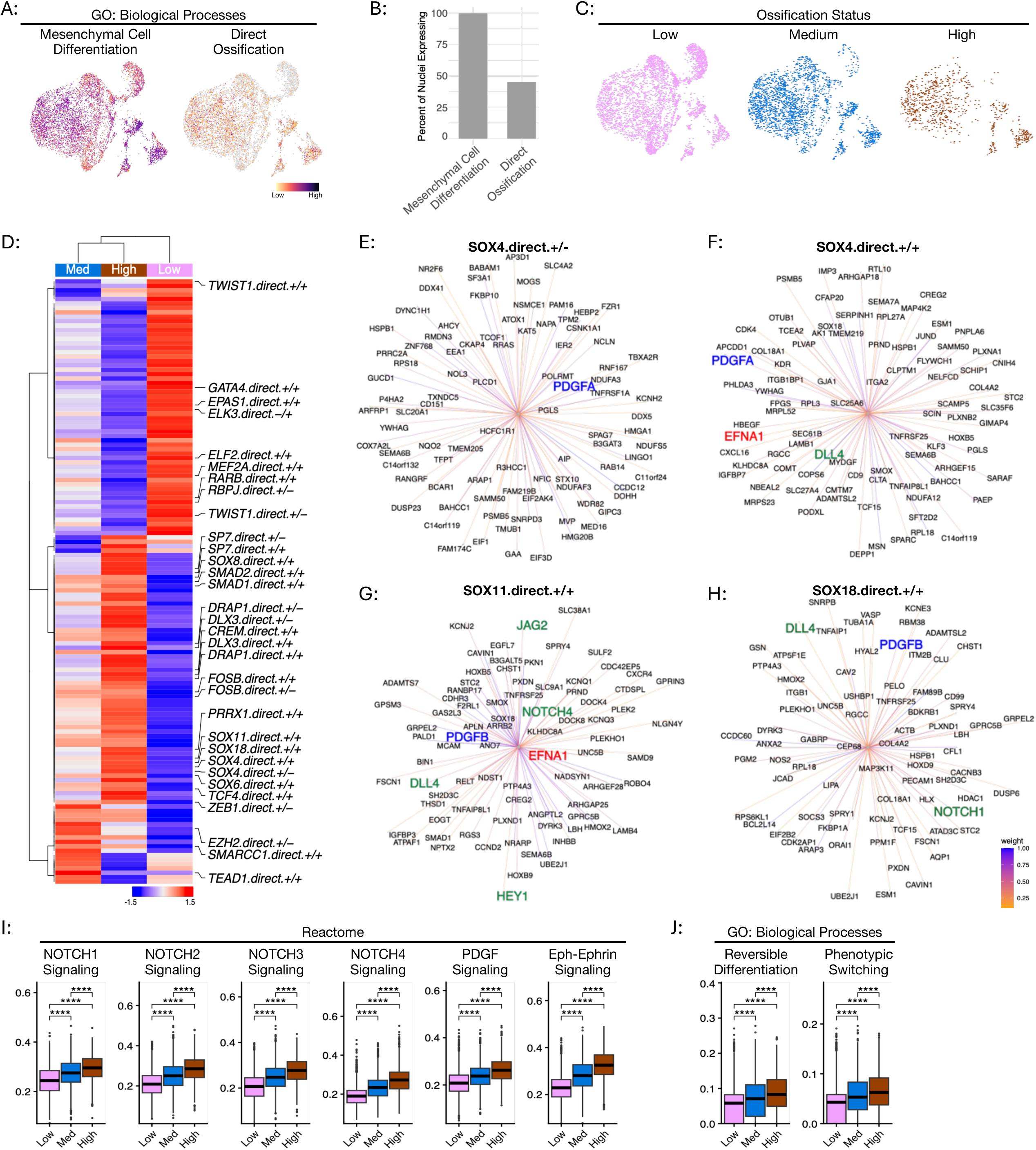
Identification and characterization of osteogenic transdifferentiation of diploid endothelial cells. (A) Gene set enrichment analysis scores for the “Mesenchymal Cell Differentiation” (left) and “Direct Ossification” (right) gene signatures displayed as UMAP plots. (B) Quantification of the percentage of nuclei expressing the “Mesenchymal Cell Differentiation” and “Direct Ossification” signatures. (C) UMAP plot showing the nuclei labeled as Ossification Low, Medium, and High. (D) Heatmap showing the results of unsupervised hierarchical clustering (Spearman) of the GRNs identified in the Ossification Low, Medium, and High groups of endothelial cell nuclei. Color bar represents z-scaled AUC values (blue = -1.5, red = +1.5) The signs following the GRN name reports the inferred activity: +/+ chromatin at locus is accessible and the target gene is expressed; +/- chromatin at locus is accessible and the target gene is not expressed; -/+ chromatin at locus is not accessible and the target gene is expressed; -/- chromatin at locus is not accessible and the target gene is not expressed. (E-H) Top target genes for the (E-F) SOX4-, (G) SOX11-, and (H) SOX18-driven GRNs. Genes of interest are highlighted in blue (PDGF signaling), red (Ephrin signaling), and green (Notch signaling). Colored lines indicate relative weight/importance as reported by triplet score in SCENIC+. (I-J) Box plots of gene set enrichment scores according to Ossification Status. (I) Reactome pathways and (J) Gene Ontology: Biological Processes signatures used in this analysis are listed above each plot. Enrichment scores are plotted on the y-axis. Asterisks indicate statistically significant differences between the groups are determined by Wilcoxon signed rank test with Benjamini-Hochberg correction post-hoc test. P<0.001 considered statistically significant.

Next, we investigated the gene regulatory networks associated with the ossification status of diploid endothelial cells. Consistent with an osteogenic program, GRNs driven by SP7, a master regulator of osteogenesis (36), were enriched in the Ossification High group and targeted genes associated with osteoblast and/or bone function including *PTH1R, B4GALTN3,* and *MESD* (**Figure 6D; Supplemental Figure 11**) (37–39). However, experimental evidence shows that SP7 alone is necessary but not sufficient for osteoblastic conversion (40). Additional GRNs enriched in the Ossification High group included those driven by transcription factors upregulated during osteogenic differentiation of human induced pluripotent stem cells such as CREB3L1, DRAP1, DLX3, FOS, MAZ, and PRRX1 (41) (**Figure 6D; Supplemental Figure 12**). Of the GRNs differentially enriched in the Ossification High group, four were driven by SOX family members SOX4, SOX11, and SOX18. Because various SOX genes often regulate stemness and lineage decisions, we used Cytotrace2 to infer the relative developmental potency/stemness of the diploid endothelial cell nuclei (42). Although there were statistically significant differences in the potency score between the diploid endothelial cells according to their ossification status, all three groups showed similar potency and were primarily classified as “Differentiated” (**Supplemental Figure 13**). These data suggest that the diploid endothelial cells undergo partial transdifferentiation towards an osteoblastic-like state rather than dedifferentiation or reprogramming followed by lineage switching.

To further explore the SOX4, SOX11, and SOX14-driven GRNs in the Ossification High group, we examined the top 50 target genes within each of these networks. This analysis repeatedly highlighted components of the Notch, PDGF, and Ephrin signaling pathways as target genes (**Figure 6E-H**), all of which have noted roles in osteogenesis. Pathway-level gene set enrichment analysis using the Reactome database further supported a statistically significant increase in Notch, PDGF, and Ephrin signaling programs in the Ossification High group (**Figure 6I**). Similar to the gene set enrichment analysis results obtained from the osteosarcoma vascular mimics, we observed statistically significant increases in the “Reversible Differentiation” and “Phenotypic Switching” signatures in the Ossification High group (**Figure 6J**). Collectively, these results indicate that a subset of diploid endothelial cells responds to osteoinductive cues within the metastatic osteosarcoma microenvironment by expressing osteogenic transcriptional programs.

### Lung microvascular endothelial cells express osteogenic transcriptional programs in response to osteosarcoma-derived factors *in vitro*

To test whether osteosarcoma cells can directly induce the expression of osteoblastic transcriptional programs *in vitro*, we performed indirect co-culture assays using a modified transwell system (**Figure 7A**). Patient-derived osteosarcoma cell lines OS186 and OS742 were grown on 3D scaffolds within transwell inserts and maintained in osteogenic growth media (OGM). Inserts were added to monolayers of primary human lung microvascular endothelial cells and cultured in OGM for the duration of the experiment. Endothelial cells cultured alone in endothelial growth media (EGM) or in OGM served as controls.

**Figure 7:**
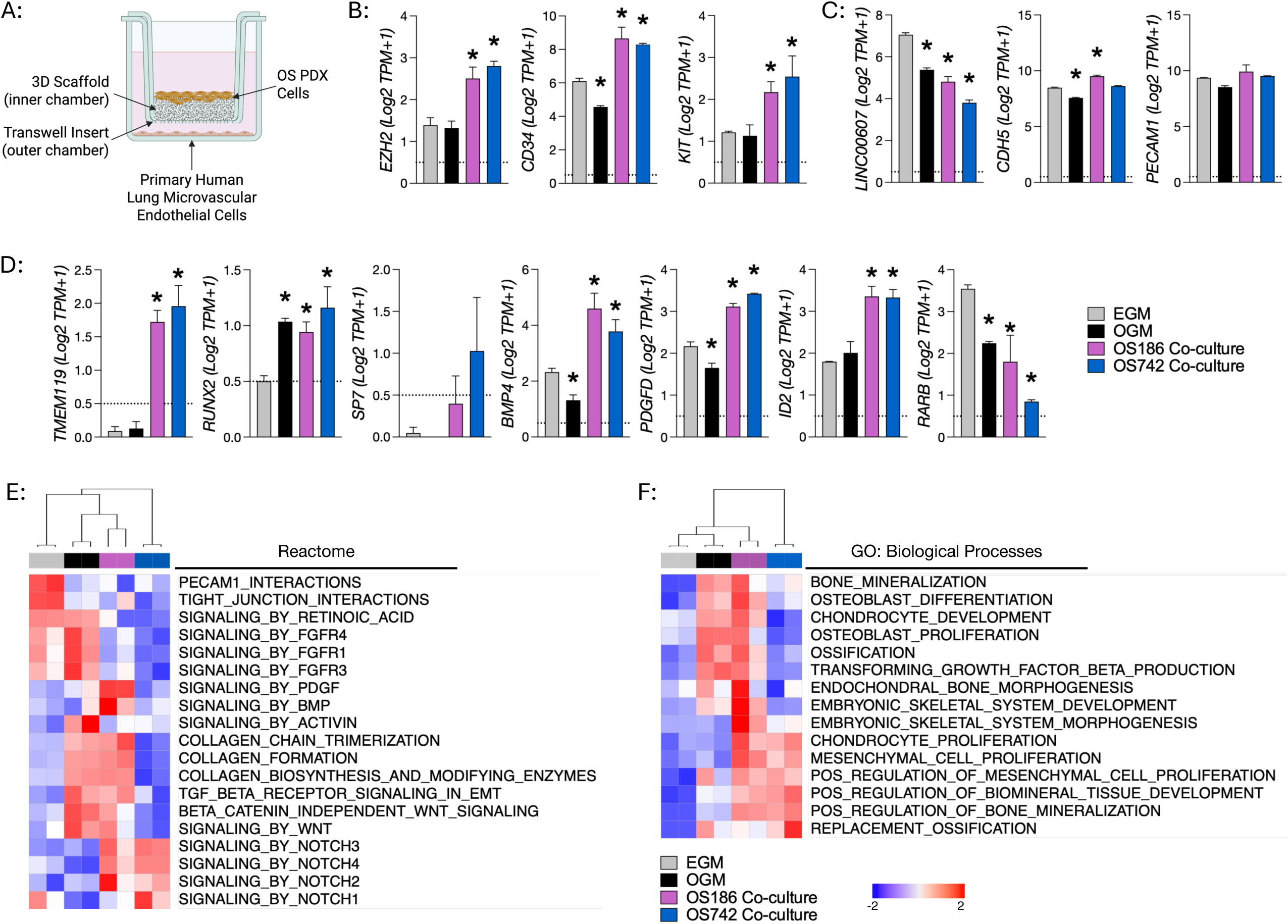
Tumor-derived factors educate lung microvascular endothelial cells to transition towards an osteoblast-like cell state *in vitro.* (A) Schematic of modified transwell co-culture assay. Image created with BioRender. (B-D) Histograms of bulk RNA-seq data. Log2 TPM+1 values plotted on the y-axis. The dotted horizontal line denotes the noise threshold, above which genes are considered reliably quantified. Asterisks indicate statistically significant differences (p<0.05) between the experimental groups and the EGM-only controls as determined by One-way ANOVA with Dunnett’s multiple comparisons test. (E-F) Heatmaps showing the results of unsupervised clustering of ssGSEA results (Pearson correlation). Colors above each column indicate sample grouping. Color bar represents z-scaled enrichment values.

Bulk RNA-seq showed that co-cultured endothelial cells upregulated *EZH2*, *CD34*, and *KIT,* consistent with acquisition of more primitive/stem-like features (**Figure 7B**). *LINC00607* is an endothelial cell enriched long non-coding RNA (43, 44). Our results show that *LINC00607* expression decreased in response to osteogenic conditions and tumor-derived factors (**Figure 7C**). In contrast, endothelial markers such as *CDH5* and *PECAM1* showed only modest changes, despite statistical significance driven by low variability (**Figure 7C**). The co-cultured primary lung microvascular endothelial cells also significantly upregulated multiple pro-osteogenic genes, including *TMEM119, RUNX2*, *BMP4, PDGFD,* and *ID2* (**Figure 7D**) (45–51). Activation of retinoic acid receptor signaling has been reported to antagonize osteogenic differentiation (52, 53). Exposure of lung microvascular endothelial cells to osteogenic medium (OGM) or osteosarcoma-derived factors significantly downregulated the expression of *RARB*, a retinoic acid–responsive nuclear hormone receptor. Notably, with few exceptions, the magnitude of gene expression changes was greater in endothelial cells co-cultured with osteosarcoma cells than in endothelial cells cultured in OGM alone. Principal component analysis of the bulk RNA-sequencing data demonstrated that each experimental condition was transcriptionally distinct from control endothelial cells cultured in EGM alone, with the largest source of variation driven by endothelial cells co-cultured with osteosarcoma cells (**Supplemental Figure 14**). These findings suggest that the observed transcriptional response is not solely attributable to exposing lung microvascular endothelial cells to OGM but rather due to osteosarcoma-derived factors *in vitro*.

To better contextualize these transcriptional changes, we performed single-sample gene set enrichment analysis. Multiple osteogenic biological processes were enriched in endothelial cells co-cultured with osteosarcoma cells and in endothelial cells cultured in OGM (**Figure 7E–F**). Analysis of the Reactome database revealed marked enrichment of Notch2, Notch3, and Notch4 signaling in endothelial cells co-cultured with either OS186 or OS742 osteosarcoma cells. In contrast, gene sets associated with canonical endothelial function, including “PECAM1 Interactions” and “Tight Junction Interactions”, were most highly enriched in endothelial cells cultured in EGM and decreased following exposure to osteogenic or tumor-derived stimuli (**Figure 7E**). Similarly, retinoic acid receptor signaling, which inhibits endothelial-to-osteoblast conversion, was also reduced in endothelial cells co-cultured with osteosarcoma cells. FGFR signaling, which antagonizes EndMT, was enriched in endothelial cells cultured in EGM and OGM but was diminished in endothelial cells co-cultured with either osteosarcoma cell line (21) (**Figure 7E**). Analysis of Gene Ontology Biological Process signatures further demonstrated enrichment of multiple osteogenic processes in endothelial cells cultured in OGM and in endothelial cells co-cultured with osteosarcoma cells, consistent with a shift away from a canonical endothelial transcriptional state and toward an osteoblast-associated program (**Figure 7F**). Together, these findings indicate that primary human lung microvascular endothelial cells are susceptible to osteosarcoma-derived factors and respond by expressing osteoblast-associated transcriptional programs.

### In vivo validation of endothelial-to-osteoblast conversion in syngeneic osteosarcoma models

While powerful and informative in its simplicity to evaluate direct transcriptional responses to stimuli, our *in vitro* co-culture assay lacks the cellular, molecular, and structural complexity to fully evaluate the response of endothelial cells to microenvironmental cues in metastatic osteosarcoma. As such, these assays are limited in their ability to capture the full spectrum of endothelial responses to tumor-derived and microenvironmental cues *in vivo*. To more accurately investigate endothelial cells in the metastatic osteosarcoma microenvironment, we injected AXT murine osteosarcoma cells paratibially into syngeneic tamoxifen-treated CDH5-CreER^T2^;Rosa26-tdTomato lineage tracing mice. The mice were treated with tamoxifen prior to receiving tumor cell injections to ensure that CDH5-expressing endothelial cells were genetically tagged by the tdTomato reporter. We performed immunohistochemistry on primary tumors and spontaneous pulmonary metastases using a monoclonal antibody against tdTomato to identify CDH5^+^ endothelial-lineage cells (**Figure 8A**). As expected, tdTomato-positive cells with vascular localization were observed in both primary and metastatic tumors, as well as in non-diseased quadriceps, bone marrow, and lung, consistent with endothelial labeling (**Figure 8B-C**). We also detected tdTomato immunoreactivity in cells embedded within osteoid matrix and not associated with vessels (**Figure 8B-C**). These osteoid-associated tdTomato-positive cells morphologically resembled osteoblasts rather than typical endothelial cells. Additionally, these tdTomato-positive cells were found in osteoid-containing regions of the tumor lesions and were absent from highly cellular regions with little or no osteoid (**Figure 8B-C**). Together, these lineage tracing data provide direct evidence that osteogenic cues within the osteosarcoma microenvironment can drive endothelial cells to acquire an osteoblast-like state, functionally supporting the computational and *in vitro* findings.

**Figure 8:**
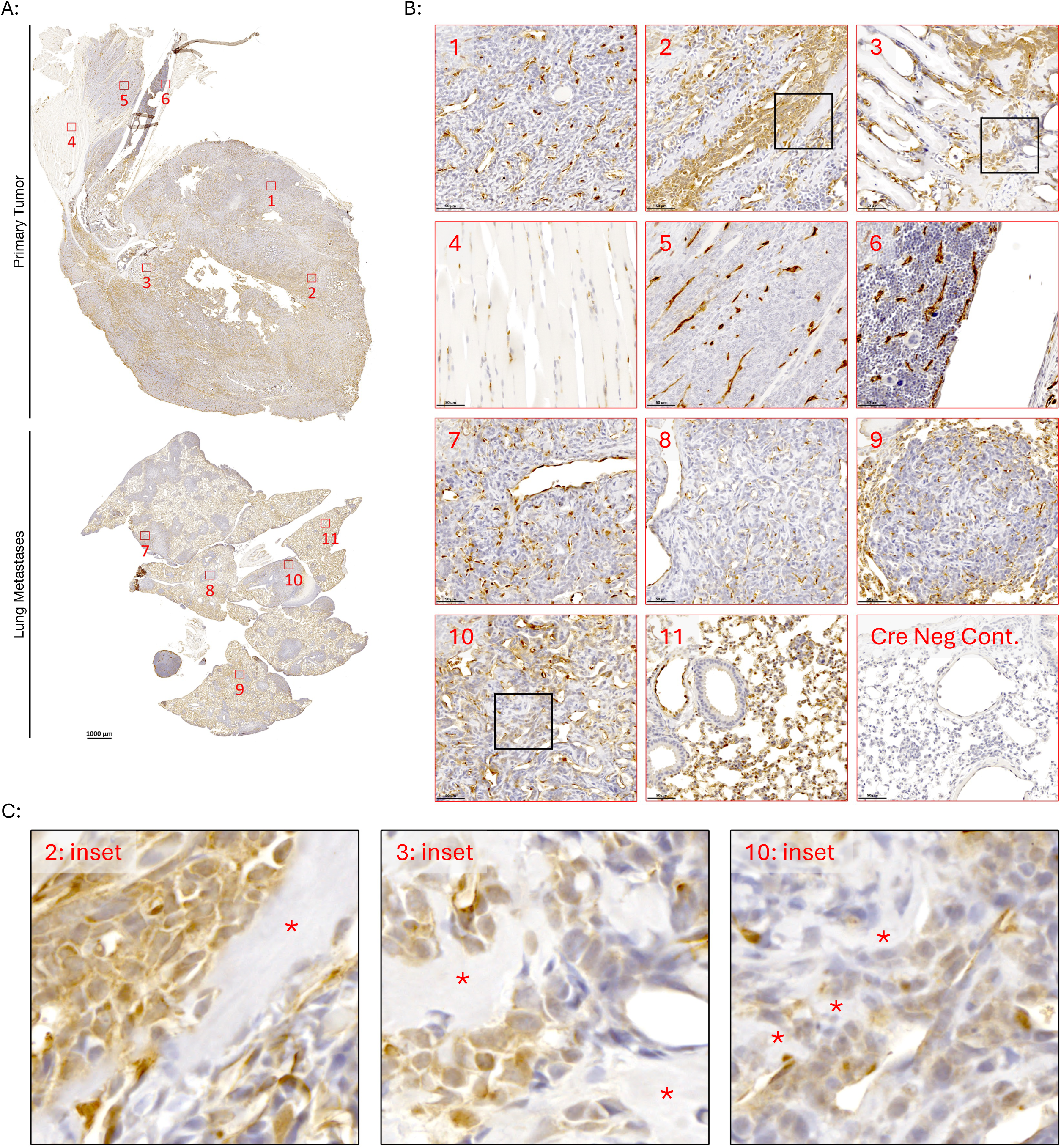
Lineage traced CDH5+ cells are embedded in osteoid-rich regions of primary and metastatic syngeneic murine osteosarcoma tumors. Immunohistochemical analysis of tdTomato^+^ endothelial cells in primary and metastatic AXT syngeneic murine tumors. TdTomato-positive cells are labeled brown and nuclei are counterstained with hematoxylin. (A) Low magnification microscopy images of osteosarcoma primary tumor (top) and lung metastases (bottom). Red boxes correspond to magnified regions of interest in (B). Scale bar = 1000um (B) Regions of interest from (A). ROIs: 1-primary tumor with scant osteoid; 2- primary tumor with significant osteoid; 3- primary tumor invading the epiphysis with significant osteoid; 4- non-affected quadricep muscle; 5-primary tumor with minimal osteoid invading quadriceps; 6- bone marrow; 7- lung metastasis with scant osteoid; 8- lung metastasis with scant osteoid; 9- lung metastasis with minimal osteoid; 10- lung metastasis with scant osteoid; 11- lung parenchyma; 12- lung parenchyma in Cre negative control animal. Scale bars = 50um. (C) Enlarged images from regions of interest from (B). Asterisks denote osteoid deposition. Images from a single representative animal are shown.

## DISCUSSION

The data presented here show that endothelial cells in metastatic osteosarcoma acquire transcriptional and regulatory programs consistent with a hybrid endothelial-mesenchymal cell state. Multiple studies have shown that signaling through TGF-β1, TGF-β2, TGF-β3, BMP2, BMP4, and BMP6 promotes EndMT, whereas FGF2 attenuates this process (54–59). Collectively, findings support a model in which a signaling imbalance within osteosarcoma pulmonary metastases favors EndMT and promotes the emergence of a hybrid endothelial-mesenchymal cell state. Because all endothelial nuclei profiled in this study exhibit mesenchymal features, we conclude that this hybrid cell state is a universal characteristic of intratumoral endothelial cells in metastatic osteosarcoma lesions. Importantly, endothelial cells that have undergone EndMT lose endothelial identity as evidenced by the downregulation of canonical endothelial cell marker genes (22). In contrast, endothelial cells in our dataset express both endothelial and mesenchymal markers, indicating that they have not completed EndMT and exist in an intermediate hybrid state. These data do not preclude the possibility that endothelial plasticity in metastatic osteosarcoma extends beyond the cell states defined in this current study. For example, some fibroblast populations within our dataset may indeed be derived from endothelial cells that have completed the EndMT process. Future studies aimed at shifting this balance to antagonize EndMT will deepen our understanding of osteosarcoma biology and may reveal therapeutic opportunities.

Vascular mimicry has been described in melanoma, high-grade glioma, breast cancer, lung cancer, and ovarian cancer (60). A histological study of primary human osteosarcoma documented vascular mimicry mediated by presumptive malignant cells based on differential staining (61). Our integrated strategy combining computational analyses with *in vivo* validation, provides compelling evidence of vascular mimicry in pediatric osteosarcoma pulmonary metastases. These data also underscore the power of integrating gene expression with chromatin accessibility to define cell states. Despite robust expression of endothelial marker genes and relatively uniform enrichment of endothelial transcriptional programs, the osteosarcoma vascular mimics segregated into distinct clusters based on their GRN profiles. Our analysis also identified two pseudotime lineages resulting in nuclei adopting endothelial-like or progenitor-like states. We therefore hypothesize that vascular mimicry represents a non-linear continuum of transitional states rather than a single discrete phenotype. Rigorous trajectory analyses will require both *in vitro* and *in vivo* functional studies using molecularly barcoded cells to determine how osteosarcoma cells progress along this continuum and whether individual cells can fluctuate between tumor-like, progenitor-like, and endothelial-like states, and if so, in what context.

Our *in vitro* data indicate that tube-forming capacity varies across osteosarcoma cell lines, suggesting that vascular mimicry is not a universal property of osteosarcoma cells. This interpretation is further supported by the observation that only a subset of tumor cells in our dataset were confidently identified as vascular mimics. Going forward, comprehensive genomic and epigenomic profiling of osteosarcoma cell lines, murine models, and patient specimens, integrated with complementary *in vitro* and *in vivo* functional studies will be essential to define the molecular drivers of vascular mimicry, elucidate the functional significance of these cells in osteosarcoma biology, and identify potential therapeutic vulnerabilities.

Endothelial-to-osteoblast conversion is an alternative physiological mechanism that contributes to bone formation and maintenance (62, 63). This process has also been observed in pathological contexts, including ectopic ossification, prostate cancer metastatic to bone, and now metastatic osteosarcoma (40, 53, 62–65). In our data, most endothelial nuclei from the liver metastasis were labeled Ossification Low. Given the substantial heterogeneity of endothelial populations within and across tissue sites, it will be important to determine whether tissue-specific endothelial subtypes are preferentially susceptible to osteoblastic differentiation (66–68). Notably, while our data suggest that all endothelial cells exist in a hybrid endothelial-mesenchymal cell state, approximately ∼40% demonstrated enrichment in the Direct Ossification signature. We therefore propose that this hybrid cell state functions as a priming event, enabling a subset of endothelial cells to further transition towards an osteoblast-like cell. The precise cues, whether cell-autonomous and/or microenvironmental, that permit these primed endothelial cells to advance toward osteoblastic differentiation in a malignant osteogenic environment remain unknown.

Osteosarcoma most commonly arises in the metaphysis of long bones, particularly within trabecular bone. During long bone growth, coinciding with the peak incidence of osteosarcoma, metaphyseal endothelial cells actively proliferate to support osteoblast development and lineage-tracing studies have shown that metaphyseal endothelial cells can undergo osteoblastic transdifferentiation (62). Together, these observations raise a provocative hypothesis: during periods of rapid long bone growth, proliferating and phenotypically plastic metaphyseal endothelial cells may be particularly susceptible to oncogenic insults and could represent a potential cell of origin. Notably, precedent exists for transdifferentiated endothelial cells serving as candidate cells of origin in rhabdomyosarcoma (69–72).

An expanding body of work highlights endothelial cells as active regulators of the tumor microenvironment, capable of shaping anti-tumor immunity through multiple mechanisms (73). As the first cellular barrier encountered by circulating lymphocytes, the endothelium plays a decisive role in determining whether immune cells can arrest, adhere, and ultimately extravasate into diseased tissues, processes that depend on a functionally intact endothelial program (74, 75). Consistent with prior reports of lymphocyte exclusion in osteosarcoma, T cells comprised <1% of all annotated cells in our dataset (7, 9, 14, 76). Further mechanistic investigations are needed to determine how the vasculature of metastatic osteosarcoma impacts lymphocyte extravasation and will be essential for defining the contribution of vascular mimicry and endothelial plasticity to immune exclusion. By focusing on the tumor vasculature, these studies may identify strategies to convert metastatic osteosarcoma from an immune-excluded to an immune-permissive tissue state, with important implications for improving the efficacy of adoptive cell therapy in this disease context.

The endothelial profiling study presented here has three major limitations. First, we were unable to obtain endothelial cells from non-diseased pediatric lung tissue to serve as a normal control for snMulti-ome sequencing. When appropriate, we leveraged scRNA-seq data from the Tabula Sapiens dataset and restricted analyses to non-diseased tissues. Second, although our lineage-tracing experiments provide definitive evidence that endothelial cells demonstrate osteoblastic properties in the AXT syngeneic osteosarcoma model, we could not determine whether these cells originated from circulating endothelial cells, bone marrow–derived endothelial precursor cells (EPCs), tissue-resident endothelial cells, or tissue-resident EPCs (77). More advanced lineage-tracing strategies will be required to resolve this *in vivo*. Finally, osteosarcoma is a rare disease, and serial sampling in pediatric patients is not permissible. As a result, our data represent a single timepoint within a complex and evolving disease process. To more accurately model vascular mimicry and endothelial plasticity, longitudinal profiling will be needed using appropriate experimental systems.

In conclusion, this multi-omic characterization of metastatic osteosarcoma endothelial cells reveals new dimensions of osteosarcoma biology and raises several important questions. Which tumor-derived factors are necessary and sufficient to drive endothelial cell state transitions towards an osteoblast-like state, are there additional microenvironmental sources of these factors, and are these factors actionable? Osteosarcoma comprises multiple histologic subtypes, including osteoblastic, chondroblastic, fibroblastic, and telangiectatic variants (78). Does the propensity for endothelial plasticity, and the resulting cellular state, vary across these subtypes? Because osteosarcoma is defined by malignant osteoid production, our findings also raise a fundamental question: What fraction of tumor-associated osteoid is produced by malignant tumor cells versus osteoblast-like endothelial cells, and does this distinction carry prognostic or therapeutic significance? Together, these unresolved questions provide a roadmap for future studies focused on the translational significance of the metastatic osteosarcoma vasculature.

## METHODS

Detailed descriptions and associated references for all computational methods and mass spectrometry analysis are provided in the **Supplemental Methods**. The patient specimen characteristics, experimental reagents, and software packages used during this study are listed in **Supplemental Tables 1-5**.

### Patient specimens

Archived frozen specimens from pediatric patients with clinically confirmed metastatic osteosarcoma were obtained from the Department of Pathology at Children’s Hospital Los Angeles under Institutional Review Board protocol CCI12-00224. Clinical attributes of the specimens are listed in **Supplementary Table 1**.

### Nuclei isolation, capture, library preparation, and sequencing

Nuclei were isolated from archived frozen metastatic osteosarcoma specimens as described in **Supplemental Methods**. The nuclei were resuspended to a concentration of 4,000-5,000 nuclei/μl and partitioned into gel beads using the Chromium Controller (10X Genomics) in accordance with the 10X Genomics User Guide (10X Genomics #CG000338). The resultant barcoded libraries were sequenced on a NovaSeq6000 (Illumina) at a depth of 50,000 reads per nucleus using the following parameters: GEX Library - 28|10|10|90, R1|I1|I2|R2 and ATAC library - 50|8|24|49, R1|I1|I2|R2).

### Data processing

Demultiplexing and alignment of the BCL files were performed using the cellranger-arc pipeline (RRID:SCR_023897). The count matrix files generated for each sample were used for downstream data analyses. The raw gene expression and chromatin accessibility data was loaded into Seurat (RRID:SCR_016341) and Signac (RRID:SCR_021158) for additional pre-processing, including filtering, normalization, dimensionality reduction, batch correction, clustering. To mitigate the effects of ambient RNA contamination, raw counts of the RNA assay were corrected using SoupX (RRID:SCR_019193), with a fixed contamination rate of 0.20, producing an adjusted gene expression matrix. Low quality nuclei and multiplets were removed from the datasets using Scrublet (RRID:SCR_018098). The RNA and ATAC assays were processed separately and then integrated using the weighted nearest neighbor (WNN) workflow in Seurat.

### Cell type annotations

ScType (RRID:SCR_026634) was used to annotate the data using the marker genes listed in **Supplemental Table 4**. Each annotation was further refined in an iterative fashion by subclustering and marker gene analysis. The top 250 marker genes for each subcluster were submitted to the Enrichr online database and queried against multiple cell type databases to determine the cell type. Subclusters were re-annotated using the corresponding cell type when consensus was reached in two or more databases. Clusters with small numbers of non-informative nuclei were removed from the dataset.

### Computational analysis

Copy number analysis on the RNA assays and ATAC assays were performed with SCEVAN and epiAneuFinder (RRID:SCR_026269), respectively. SCENIC+ (RRID:SCR_026702) was used to build gene regulatory networks. Gene set enrichment analysis was performed on individual nuclei using the UCell package (RRID:SCR_027109). Transcription factor motif activity analysis was performed using Chromvar (RRID:SCR_026570). CytoTrace2 (RRID:SCR_022828) was used to assess stemness/potency.

### Data visualization

Dimensional reduction UMAP plots and ridge plots were created using Seurat (RRID:SCR_016341). The scCustomize package (RRID:SCR_024675) was used to generate feature plots and dot plots. Blended UMAP plots and box plots were created using the SeuratExtend package (RRID:SCR_026143). Bar graphs, pie charts, and stacked bar graphs were made using ggplot2 (RRID:SCR_014601). Heatmaps were generated using the ComplexHeatmap package (RRID:SCR_017270). The EnhancedVolcano package was used to create volcano plots (RRID:SCR_018931). GRN network plots were created using igraph (RRID:SCR_019225), tidygraph (RRID:SCR_027617), ggraph (RRID:SCR_021239), and ggplot2 (RRID:SCR_014601).

### Cell lines

Human patient derived osteosarcoma cell lines OS186 (RRID:CVCL-C8FT), OS052 (RRID:CVCL-C8FR), OS384 (RRID:CVCL-C8FU), and OS742 (RRID:CVCL-C8FY) were obtained from Dr. Alejandro Sweet-Cordero (33). Early passage cells were used for the experiments, and the cells were confirmed free of mycoplasma contamination. For routine culture/maintenance, the cells were grown in DMEM (Invitrogen #10566016) + 10% bovine growth serum (Cytiva #SH30541.03) + antibiotic/antimycotic (Invitrogen #15070063). Two weeks prior to experimentation, the growth conditions were switched to complete osteoblast growth media (Lonza #CC-3207). AXT murine osteosarcoma cell lines were obtained from Dr. Takatsune Shimizu (34). The cells were confirmed to be free of murine pathogens and mycoplasma. The cells were grown in IMDM (Invitrogen #31980030) + 10% fetal bovine serum (Omega Scientific #FB-02). Primary human lung microvascular endothelial cells were purchased from Lonza and cultured in complete endothelial growth media according to the manufacturers suggestion (Lonza #CC-3202).

### Tube formation assays

Tube formation assays were performed using µ-Plate 96 Well 3D plates (ibidi, Cat. No. 89646). Each inner well was coated with 10 µL of growth factor-reduced Matrigel (Corning, Cat. No. 354230) and incubated at 37°C for 30 minutes to allow gel polymerization prior to cell seeding. The osteosarcoma cell lines OS052, OS384, OS186, and OS525 were seeded at 10,000 cells per well in 70 µL of complete osteoblast growth media (Lonza #CC-3207). Plates were maintained in an IncuCyte® Live-Cell Analysis System (Sartorius), and phase-contrast images were acquired at 4× magnification at 1, 4, 8, 12, and 16 hours after cell seeding. Tube formation was quantified from images acquired at each time point using the Angiogenesis Analyzer plugin in Fiji/ImageJ. Morphometric parameters, including total tube length, number of junctions, branches, and meshes, were measured as using the Angiogenesis Analyzer software framework (79). This experiment was performed in biological triplicates.

### Endothelial-osteosarcoma co-culture assays

Human osteosarcoma patient derived cell lines were grown in complete osteoblast growth media (Lonza #CC-3207) then plated into Alvetex 3D scaffolds prepared according to the manufacturers instructions (Amsbio #AMS.AVP004-96). Osteosarcoma cells were grown on 3D scaffolds for 24 hours prior to co-culture. Human microvascular lung endothelial (HMVEC-L) cells (Lonza #CC-2527) were grown in complete endothelial cell media (Lonza #CC-3202) and plated on gelatin-coated 6-well plates (Corning #356652) at passage 1. The HMVEC-L cells were grown to confluence then starved overnight in Opti-MEM (Invitrogen #31985070). The next day, the starvation media was replaced with fresh endothelial growth media (EGM), osteoblast growth media (OGM), or OGM supplemented with 100ng/mL BMP4 (R&D Systems #S67001). Osteosarcoma-containing 3D scaffolds were placed into the top of transwell inserts (Corning #353091) and the inserts were added to endothelial cells with OGM. For each condition, 50% of the media was refreshed every two days for 10 days. On day 10, RNA was extracted from the endothelial cells using the RNA/DNA/Protein Purification Plus kit (Norgen Biotek #47700) and submitted for bulk RNA sequencing. Biological duplicates of this experiment were performed.

### Bulk RNA sequencing

Poly- A libraries were constructed using the NEBNext Ultra II Directional RNA Library Prep kit for Illumina (New England Biolabs #7760) as per the manufacturer’s protocol. Paired-end sequencing was performed on the NextSeq2000 (Illumina). Demultiplexing was performed using the DRAGEN BCL Convert (v 4.2.7) workflow and subsequently processed using the RENEE RNA sequencing analysis pipeline (v2.6.7) (https://doi.org/10.5281/zenodo.17315880). Raw paired-end FASTQ files were aligned to the human reference genome hg38 using the hg38_45 reference bundle (GENCODE release 45) with STAR v2.7.6a (RRID:SCR_004463). Gene-level quantification was performed using RSEM v1.3.3 (RRID:SCR_000262) to generate gene-level expected counts. Log2 transformed gene-level counts were used to create histograms (GraphPad Prism, RRID:SCR_002798). Single sample gene set enrichment analysis (SSGSEA) was performed using the Reactome, Gene Ontology: Biological Processes, and WikiPathways libraries from the Molecular Signatures Database (GenePattern; RRID:SCR_003201). Unsupervised hierarchical clustering of the SSGSEA enrichment scores were displayed as heatmaps (Morpheus; RRID:SCR_017386).

### In vivo studies

The *in vivo* studies were performed under ACUC protocol POB-001. CDH5-Cre^ERT2^; Rosa26-tdTomato mice were created by crossing heterozygous CDH5-Cre^ERT2^ mice with B6.Cg-Gt(ROSA)26Sor^tm(CAG-tdTomato)Hze/J^ mice (Jackson #007909; RRID:IMSR_JAX:007909) (80, 81). The resultant mice were used for the endothelial lineage tracing experiments in this study. Primers and thermocycling conditions used for genotyping are listed in **Supplemental Table 5**. Mice received an intraperitoneal injection of 20mg/mL tamoxifen (Sigma #T176-10MG) dissolved in corn oil (SelleckChem #S6701) at a dose of 75mg/kg for three consecutive days starting at postnatal day 14. Wild-type C57Bl6/J mice were purchased from Jackson Labs (Jax stock #000664; RRID:IMSR_JAX:000664) and used for the vascular mimicry study. For each study, 4-week-old mice (n=3-4) received a 50μl paratibial injection of 2×10^6^ viable AXT cells suspended in 50% growth factor-reduced Matrigel (Corning #356231) in Hanks buffered saline solution (Invitrogen #14170112). Injections were not performed if the if cells were < 95% viable as determined by trypan blue exclusion assay. Tumors were grown for 28 days at which point the mice were euthanized.

### Tissue processing

Tissues were fixed *in situ* via trans-cardiac perfusion with PBS (Invitrogen #14190144) followed by 10% neutral buffered formalin (VWR #16004-115) then excised and decalcified in Formical-4 (StatLab #NC1053842) for 5 hours at room temperature. The decalcified tissues were placed in 10% neutral buffered formalin for 16-hours at 4C, washed in Tris buffered saline, and then placed in 70% ethanol until processing. The samples were embedded in paraffin, sectioned at 5um thickness, and mounted on to positively charged glass slides.

### Immunohistochemistry (IHC)

Formalin fixed paraffin embedded (FFPE) sections were deparaffinized, rehydrated, and subjected to heat-mediated antigen retrieval in citrate buffer (Vector Laboratories #H-3300-250; RRID:AB_2336226). Endogenous phosphatases and peroxidases were quenched using BLOXALL reagent (Vector Laboratories #SP-6000-100; RRID:AB_2336257). Specimens were then blocked with 2.5% horse serum (Vector Laboratories #S-2000-20; RRID:AB_2336617) diluted in 1X Tris-buffered saline + 0.01% Tween-20 and incubated with a rabbit monoclonal antibody against tdTomato (Cell Signaling Technology # 20163; RRDI:AB_3662938) or a goat polyclonal antibody against GFP (Abcam #ab6673, RRID:AB_305643) overnight at 4°C. Color development was achieved by incubating the slides with a peroxidase-conjugated anti-rabbit secondary antibody (Vector Laboratories #MP-7401; RRID: AB_2336529) followed by 3,3’-diaminobenzidine (Vector Laboratories, SK-4105; RRID: AB_2336520) or an alkaline phosphatase-conjugated anti-goat secondary antibody (Vector Laboratories #MP-5405; RRID:AB_2828010) followed by ImmPACT Vector Red substrate (Vector Laboratories #SK-5105; RRID:AB_2336524). Images were captured on an Axioscan 7 slide scanner or an AxioObserver microscope (ZEISS).

### Mass spectrometry

Frozen tissues were lysed in T-PER reagent (Invitrogen #78510) on dry ice and supplemented with protease inhibitor cocktail at a final concentration of 2X (Invitrogen #78425). Samples were submitted to the Protein and Metabolite Characterization Core at the Frederick National Laboratory for Cancer Research for tandem mass tag (TMT) mass spectrometry. Detailed descriptions of the methods and procedures are in the **Supplemental Methods**.

## CONFLICT OF INTEREST

The authors declare no financial conflicts of interest as it pertains to the data presented in this manuscript.

## Data availability

The data deposition is currently progress. The manuscript will be updated with the dbGaP accession number once it is issued. The snMultiome data analysis pipeline is available at https://github.com/NCI-CCDI/ccrccdi5_os_snmultiome. The bulk RNA sequencing data analysis pipeline is available at https://doi.org/10.5281/zenodo.17315880.

## AUTHOR CONTRIBUTIONS

J. Burks: Performed experiments, analyzed the data. Y. Wu: Processed the raw data, performed computational analysis, wrote the manuscript. K. Bhuvaneshwar: Processed the raw data and performed computational analysis. N. Syed: Performed experiments, processed the data. C.M. Sayers: Performed experiments. D.O. Williams: Performed experiments, analyzed the data. S.V. Daulatabad: Analyzed the data, wrote the manuscript. T. Malone: Specimen acquisition, data curation. J. Galindo: Specimen acquisition, data curation. M. Mendez: Specimen acquisition, data curation. J. Cotter: Specimen acquisition, data curation. J. Pavisic: Conceptualization, analyzed the data. Y. Mukouyama: Conceptualization, resources, edited the manuscript. J.F.Shern: Conceptualization, edited the manuscript. R.N.Kaplan: Conceptualization, edited the manuscript. T.A. McEachron: Conceptualization, data analysis, wrote and edited the manuscript.

## ACKNOWLEDGMENTS

This research was supported, in part, by the Intramural Research Program of the National Institutes of Health (ZIABC012056 to TAM), the NCI Childhood Cancer Data Initiative (3P30CA008748-54S3), and Federal funds under Contract No. 75N91019D00024. The contributions of the NIH author(s) were made as part of their official duties as NIH federal employees, are in compliance with agency policy requirements, and are considered Works of the United States Government. However, the findings and conclusions presented in this paper are those of the author(s) and do not necessarily reflect the views of the NIH or the U.S. Department of Health and Human Services. The Center for Pathology Research Services at Children’s Hospital of Los Angeles and the Norris Comprehensive Cancer Center Translational Pathology Core are supported by grant 5P30CA014089-45. This work utilized the computation resources of the NIH High-Performance Computing Biowulf cluster (http://hpc.nih.gov). We thank Dr. Carol Thiele for her critical review of the manuscript. Text was refined for clarity and grammar using ChatGPT (OpenAI, HHS government version).

## Notes

### Competing Interest Statement

The authors have declared no competing interest.

