## Supplemental Figures and Figure Legends for "Integrated multi-omic analysis of pediatric metastatic osteosarcoma reveals endothelial cell plasticity and lineage infidelity"

#### SUPPLEMENTAL FIGURE LEGENDS

**Supplemental Figure 1: Clustering and annotation of the snMulti-ome dataset.** (A) UMAPs of clustering results performed on the RNA data (left), ATAC data (middle), and WNN integrated data (right). (B) UMAP of the WNN integrated data partitioned by annotation. (C) Stacked bar graph showing the relative cellular composition of each specimen.

### Supplemental Figure 1

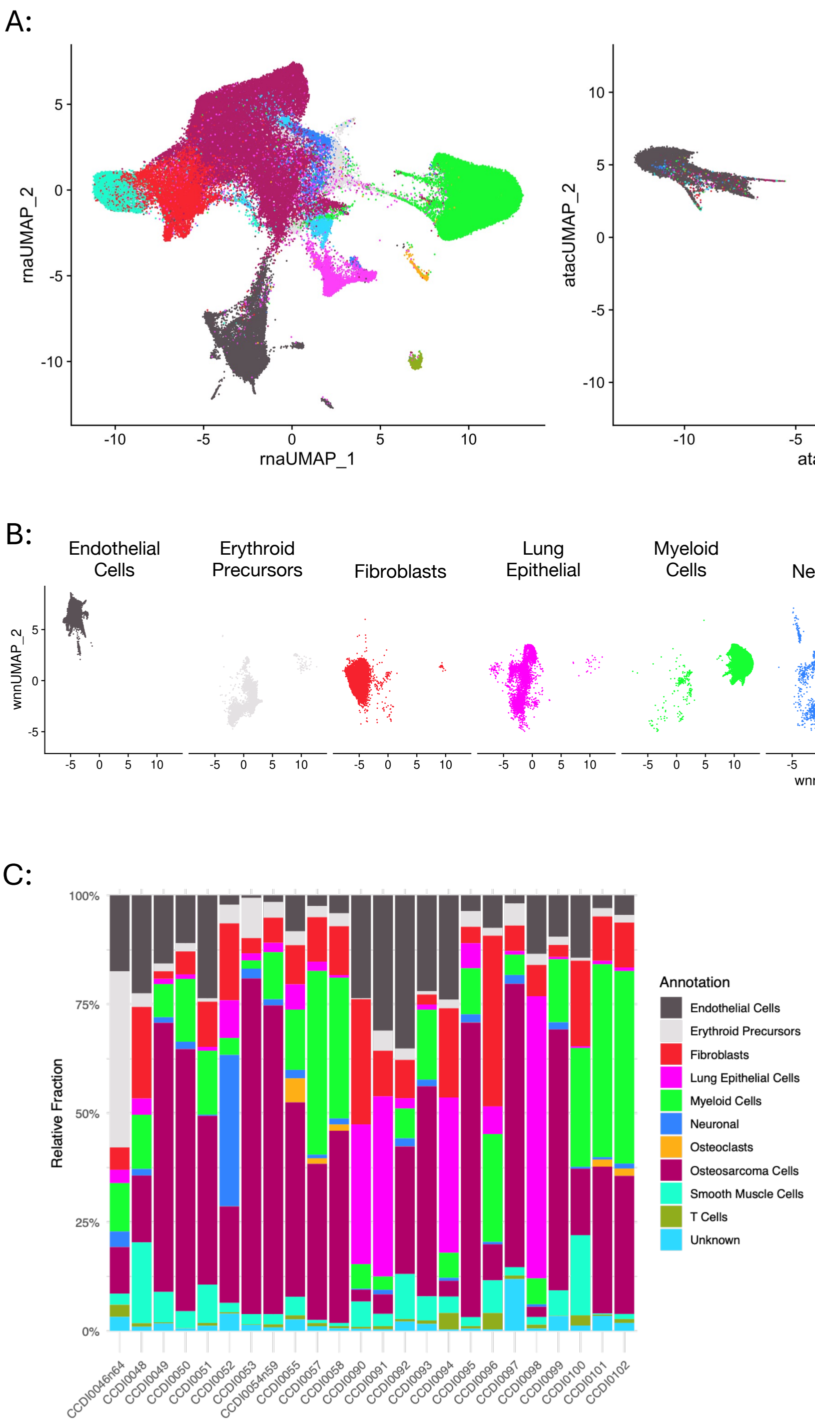

**Supplemental Figure 2: Endothelial cell marker genes.** Clustered dot plot showing the expression of endothelial cell marker genes in relation to other cell types annotated within the dataset.

#### Supplemental Figure 2

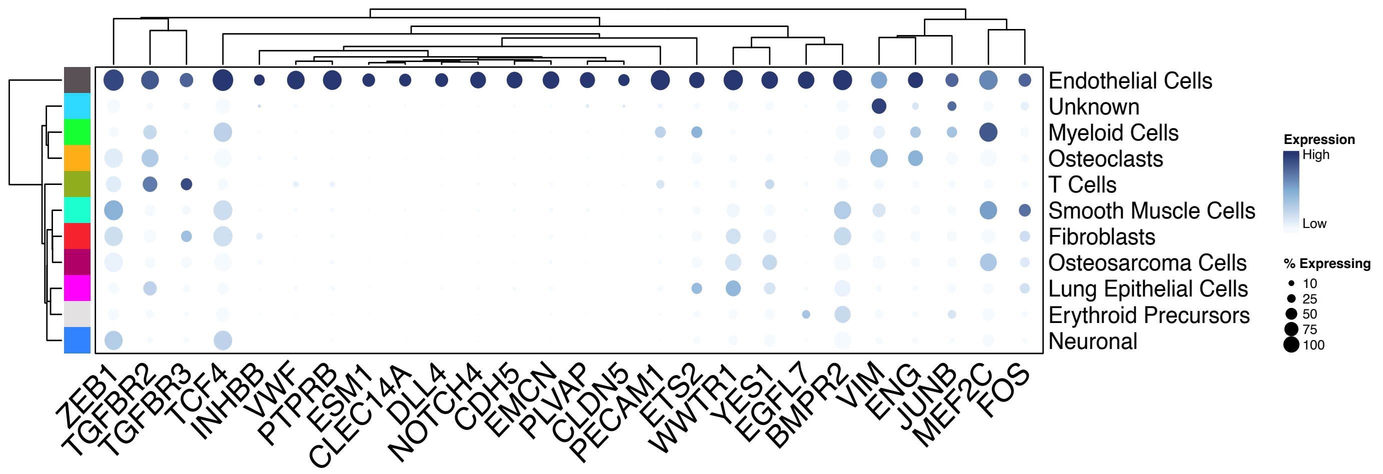

**Supplemental Figure 3**: **Composition of the subsetted endothelial cell object.** (A) UMAP plot showing the specimen identity of individual nuclei to highlight the absence of batch effect. (B) Stacked bar graph showing the relative proportion of the different endothelial cell clusters in each specimen.

### Supplemental Figure 3

A:

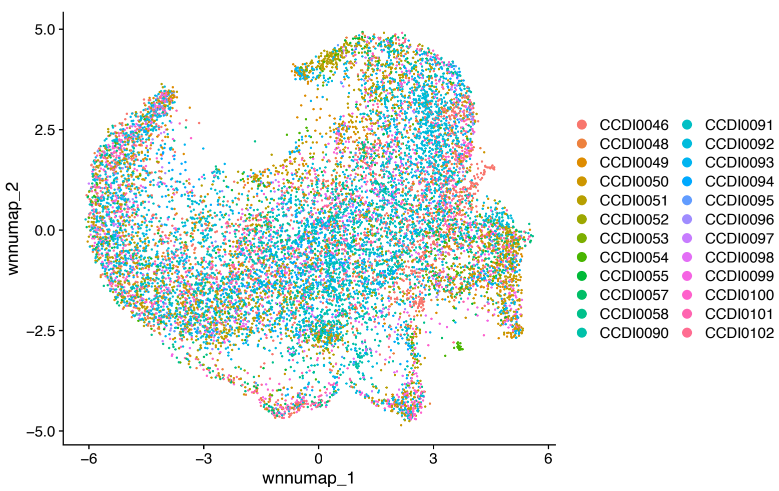


B:

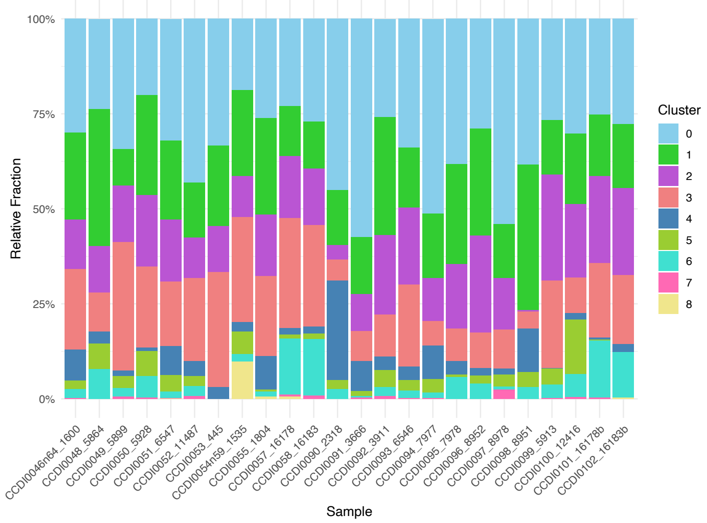

**Supplementary Figure 4. Comparative analysis of endothelial cell-state similarity using RNA, ATAC, and integrated WNN modalities.** Heatmaps comparing the similarity among endothelial cell states. (A) RNA-based similarity calculated as the Spearman correlation between average gene expression profiles for each cell state. (B) ATAC-based similarity calculated from Euclidean distances between cell-state centroids in latent semantic indexing (LSI) space. (C) Integrated similarity based on connectivity in the weighted nearest neighbor (WNN) graph. Diagonal values were normalized to 1 and represent maximal within-cell-state connectivity, whereas off-diagonal values indicate the relative similarity between cell states in the integrated RNA–ATAC neighborhood graph. In all heatmaps, rows and columns were hierarchically clustered using Euclidean distance and complete linkage.

**Supplemental Figure 4:**

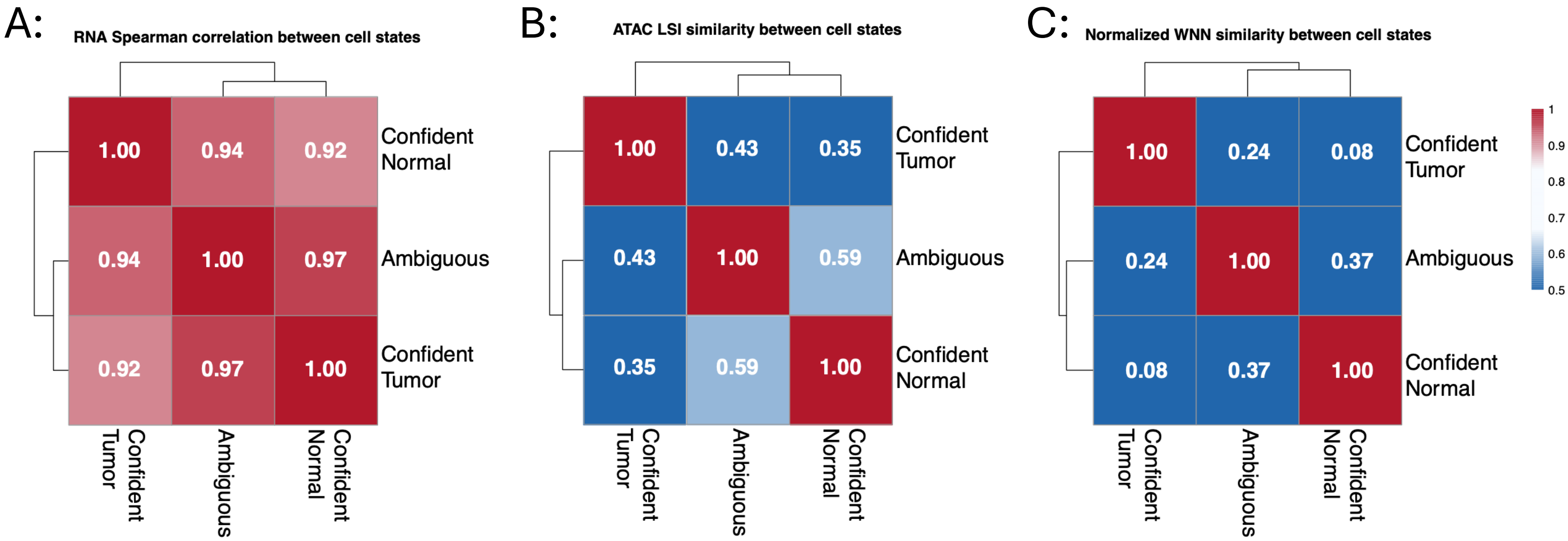

**Supplemental Figure 5: Results of epiAneuFinder and SCEVAN copy number analysis.** (A) The relative proportion of nuclei labeled as normal, tumor, or N/A using SCEVAN. (B) The relative proportion of nuclei (right) of labeled as normal or tumor using epiAneuFinder. Filter indicates the nuclei that were filtered out due to insufficient number of high-quality fragments. (C) The relative proportion nuclei labeled as Ambiguous, Confident Normal, or Confident Tumor after integration of the SCEVAN and epiAneuFinder results. (D) Pie chart showing the distribution of final CNV calls after integration.

### Supplemental Figure 5

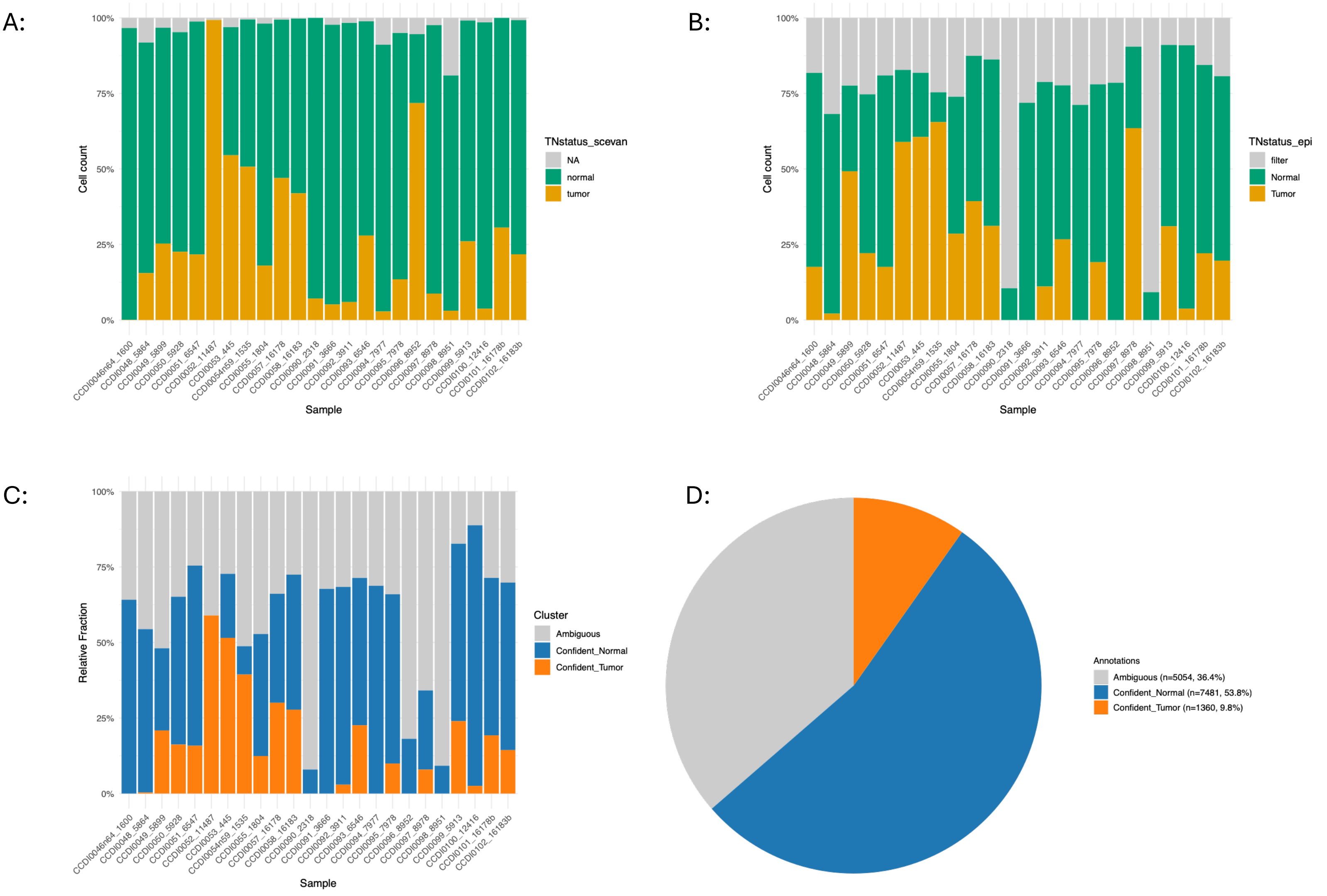

**Supplemental Figure 6**: **Composition of the diploid endothelial cell and osteosarcoma vascular mimics.** (A-B) UMAP plots showing the specimen identity of individual nuclei in the (A) diploid endothelial cell and (B) osteosarcoma vascular mimics. (C-D) UMAPs of the (C) diploid endothelial cell data and (D) osteosarcoma vascular mimics partitioned by disease site.

### Supplemental Figure 6

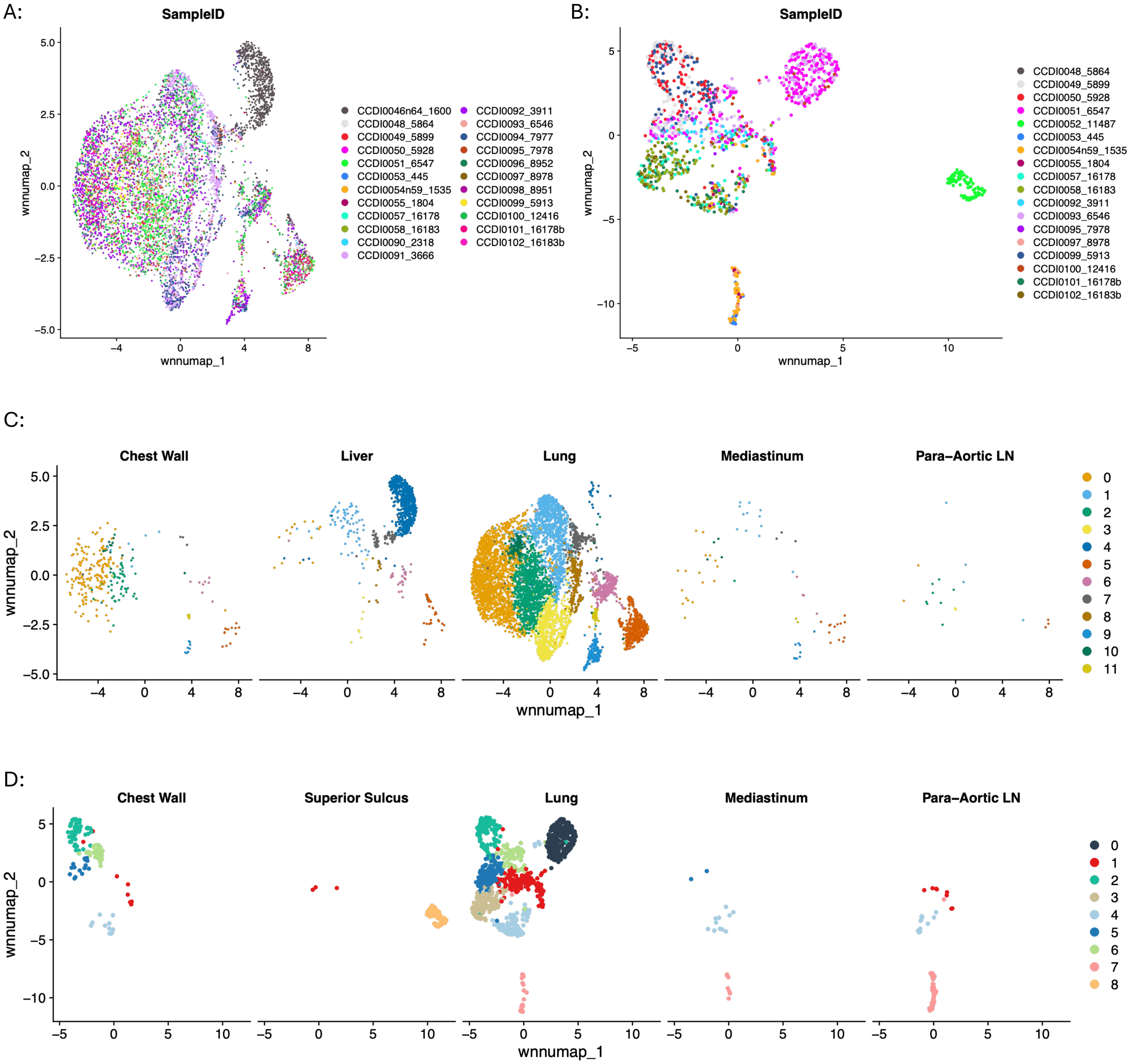

**Supplemental Figure 7: Heatmap of osteosarcoma vascular mimics GRNs.** Heatmap showing the results of unsupervised clustering (Spearman correlation) of the GRN data for the osteosarcoma vascular mimics. All GRNs are labeled to the right. The signs following the GRN name reports the inferred activity: +/+ chromatin at locus is accessible and the target gene is expressed; +/- chromatin at locus is accessible and the target gene is not expressed; -/+ chromatin at locus is not accessible and the target gene is expressed; -/- chromatin at locus is not accessible and the target gene is not expressed. Colors above each column indicate VM grouping. Color bar represents z-scaled AUC values.

### Supplemental Figure 7

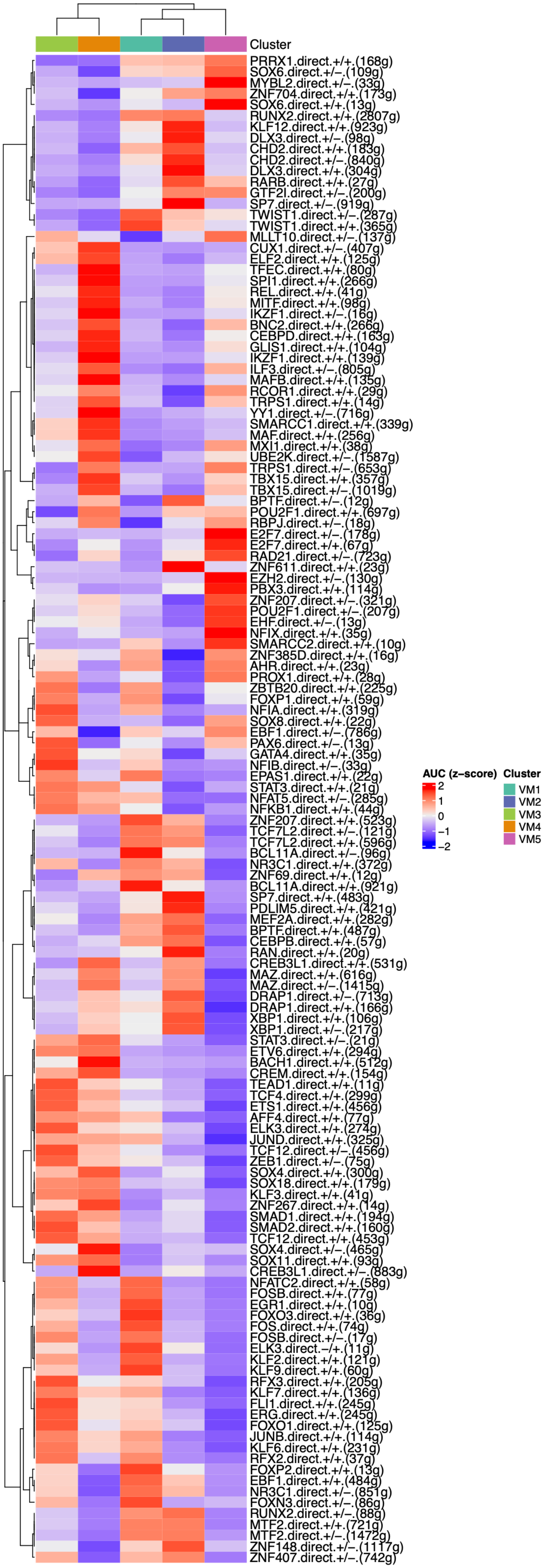

**Supplemental Figure 8: Osteogenic gene expression in vascular mimics.** Boxplots showing the expression of known osteogenic/osteoblast-related genes in each vascular mimicry (VM) grouping. Genes are listed above each plot. Gene expression values are plotted on the y-axis. Asterisks indicate statistically significant differences between the groups are determined by Wilcoxon signed rank test with Benjamini-Hochberg correction post-hoc test. $P<0.001$ considered statistically significant.

#### Supplemental Figure 8

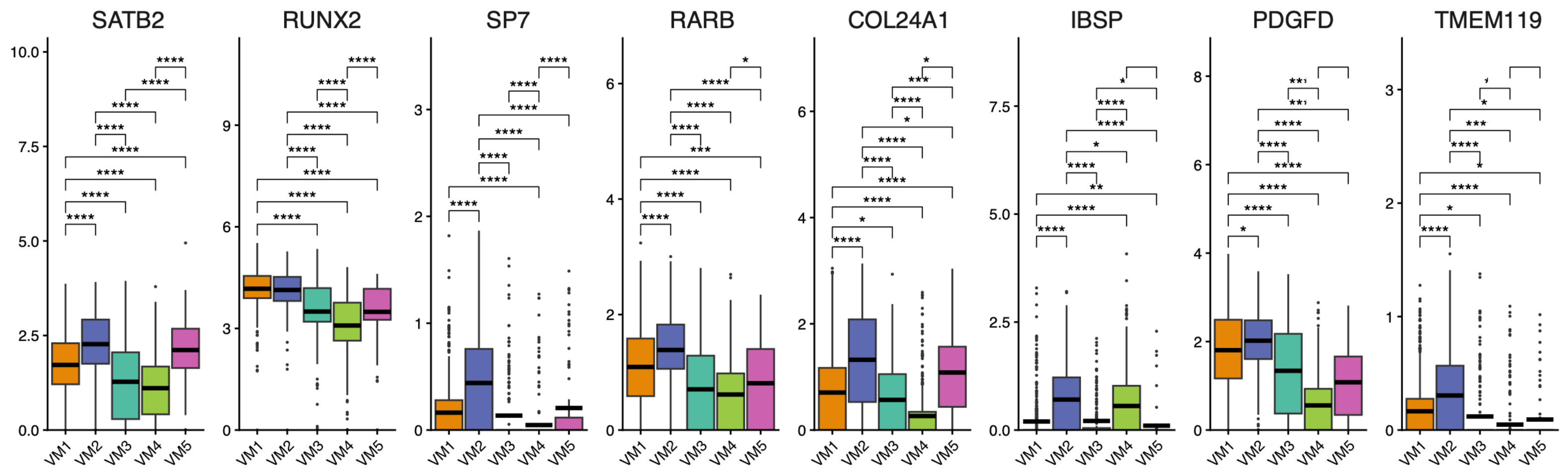

**Supplemental Figure 9: Developmental potency/stemness of osteosarcoma vascular mimics.** (A-B) UMAP plots of (A) RNA-based clustering and (B) cluster assignments after GRN-based clustering and re-annotation. (C-D) Pseudotime lineages projected on to UMAP plots showing relative ordering of nuclei in (C) Lineage 1 and (D) Lineage 2. Color bar indicates relative pseudotime. (E) Boxplots showing *PRRX1* expression. Gene expression values are plotted on the y-axis. Asterisks indicate statistically significant differences between the groups are determined by Wilcoxon signed rank test with Benjamini-Hochberg correction post-hoc test. $P<0.001$ considered statistically significant. (F) Boxplot showing the potency/stemness of the nuclei in the VM groups. Left Y-axis is the potency score. Right Y-axis is the differentiation status. Lower potency score = more differentiated. Statistical significance determined by Wilcoxon signed rank test with Benjamini-Hochberg correction post-hoc test. $P<0.001$ considered statistically significant.

#### Supplemental Figure 9

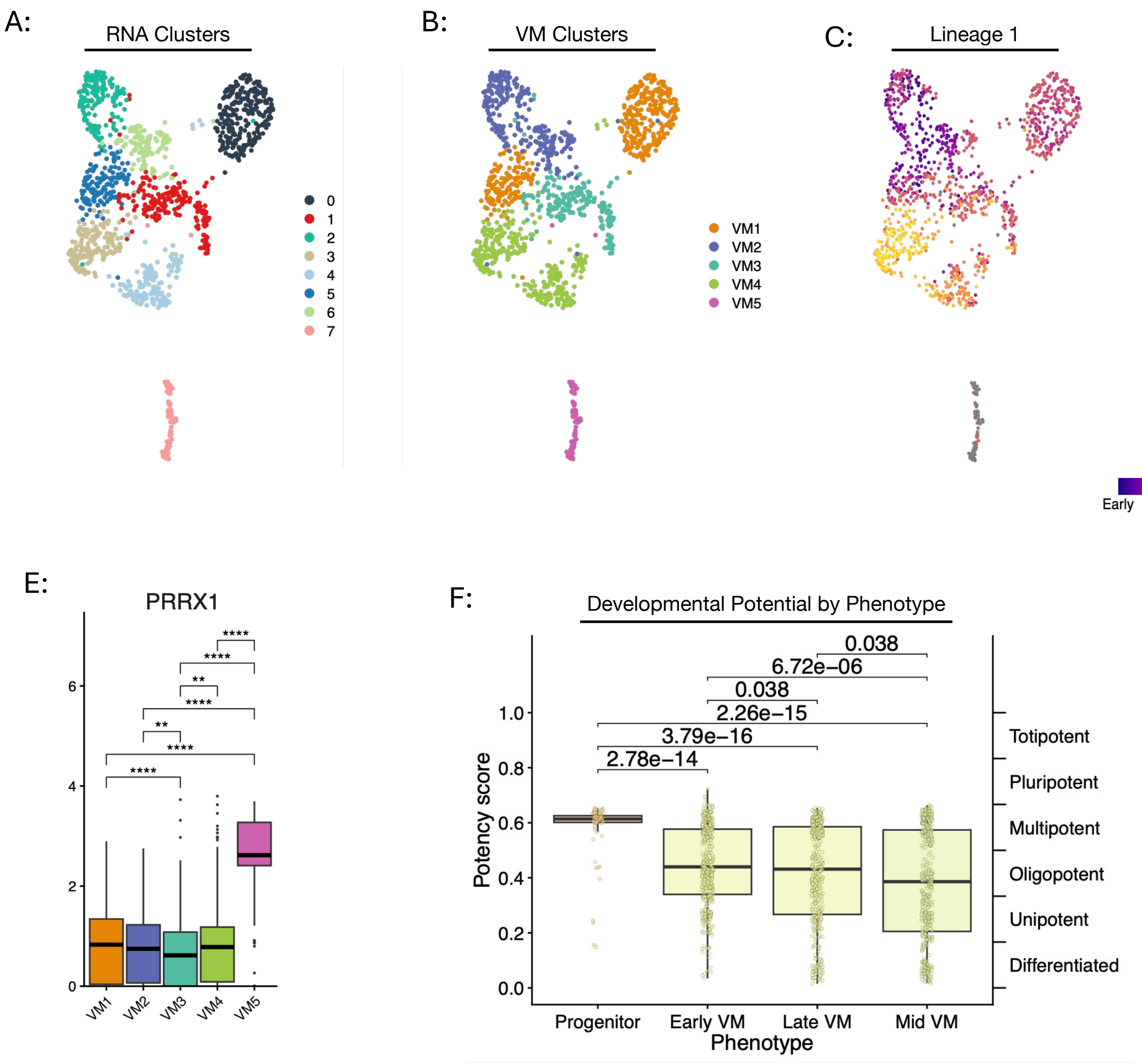

**Supplemental Figure 10: Partitioning diploid endothelial cells by ossification status.** (A) UMAP of diploid endothelial cell nuclei shown for reference. (B) Ridge plots showing the cluster-specific distribution of the "Direct Ossification" gene signature in the diploid endothelial cell nuclei. Y-axis represents the number of nuclei within the given cluster assignment colored according to UMAP shown in (A). Vertical red lines indicate thresholds for Ossification Low, Medium, and High groups. (C) Stacked bar graph showing the relative composition of each cluster in the Ossification Low, Med, and High groups.

### Supplemental Figure 10

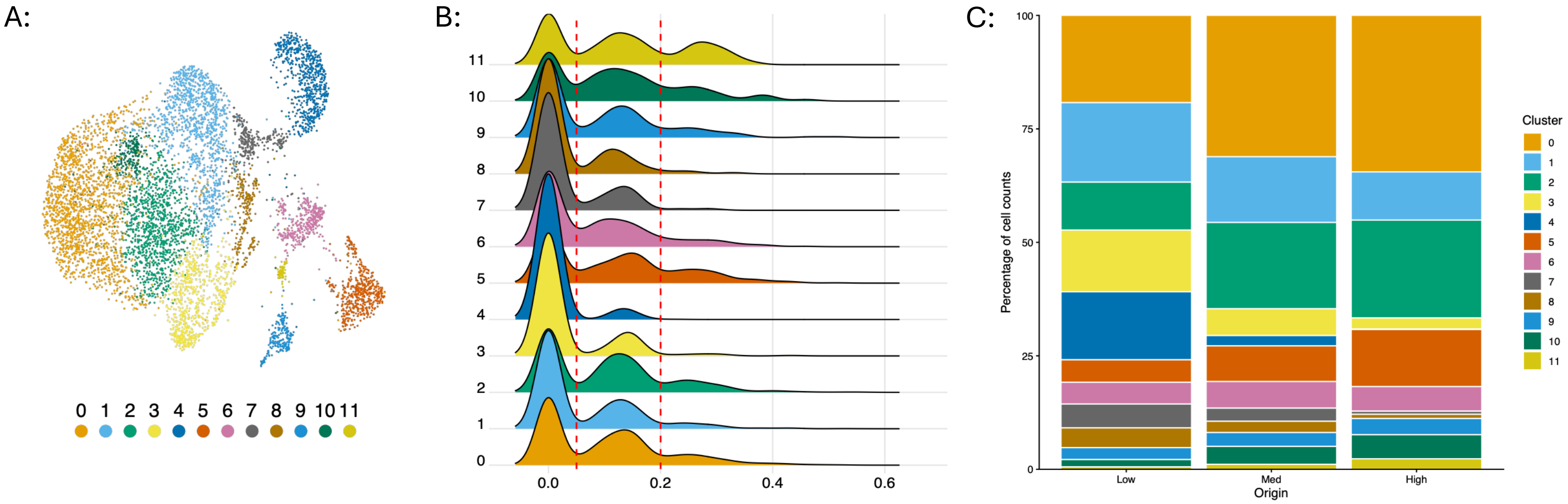

**Supplemental Figure 11: Target genes in SP7-driven GRNs in osteosarcoma VM Late nuclei.** Top target genes in activating (left) and repressive (right) SP7-driven GRNs. The signs following the GRN name reports the inferred activity: +/+ chromatin at locus is accessible and the target gene is expressed; +/- chromatin at locus is accessible and the target gene is not expressed. (E-F) Osteogenic genes of interest are highlighted in red. Colored lines indicate relative weight/importance as reported by triplet score in SCENIC+.

### Supplemental Figure 11

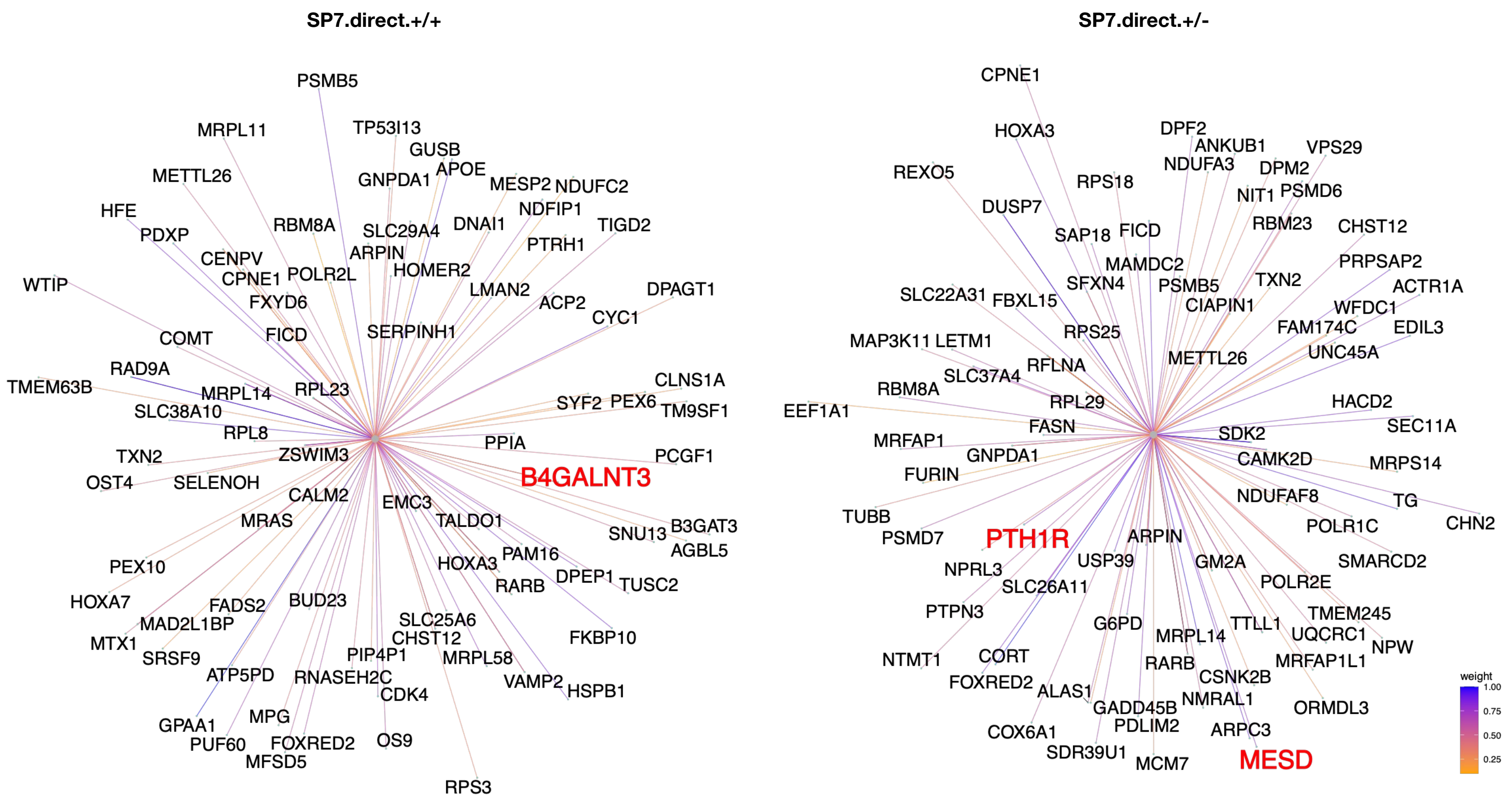

**Supplemental Figure 12: Heatmap of diploid endothelial cell GRNs.** (A) Heatmap showing the results of unsupervised clustering (Pearson correlation) of the GRN data for the Ossification Low (pink), Medium (blue), and High (brown) endothelial cell nuclei. The signs following the GRN name reports the inferred activity: +/+ chromatin at locus is accessible and the target gene is expressed; +/- chromatin at locus is accessible and the target gene is not expressed; -/+ chromatin at locus is not accessible and the target gene is expressed; -/- chromatin at locus is not accessible and the target gene is not expressed. Colors above each column indicate cluster assignment. Color bar represents z-scaled AUC values.

#### Supplemental Figure 12

A:

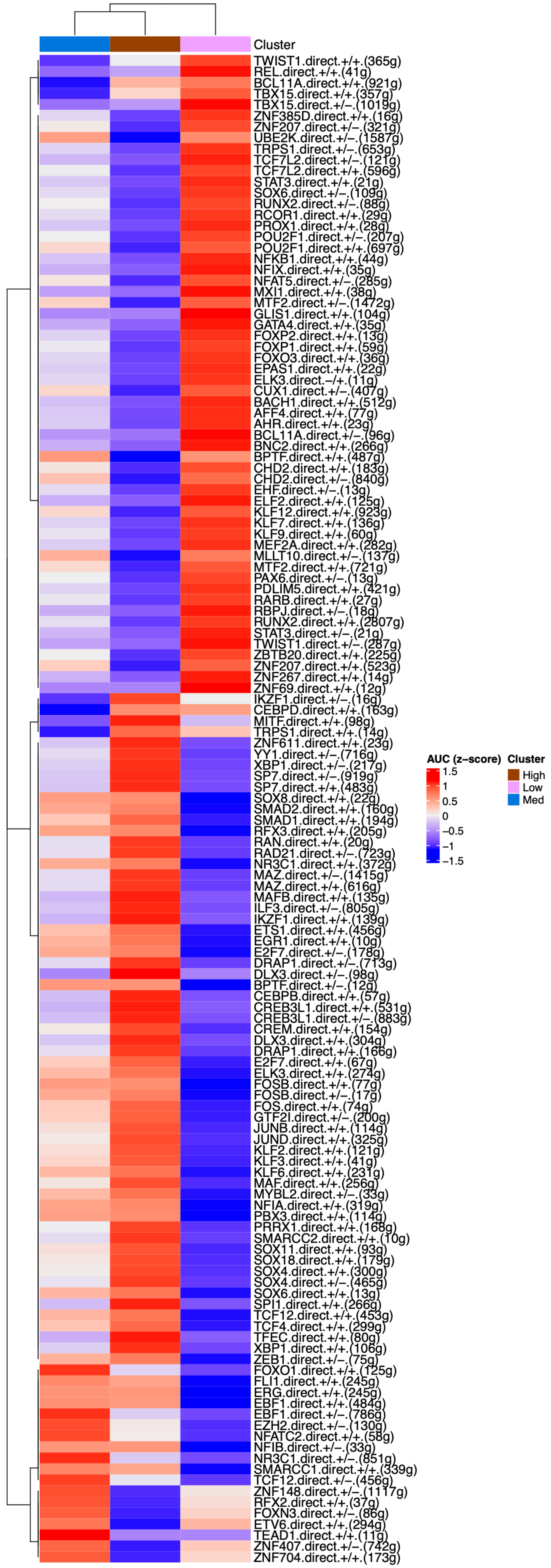

**Supplemental Figure 13: Results of CytoTrace2 Analysis.** Boxplot showing the potency/stemness of the nuclei in the Ossification Low, Medium, and High groups. Left Y-axis is the potency score. Right Y-axis is the differentiation status. Lower potency score = more differentiated. Statistical significance determined by Wilcoxon signed rank test with Benjamini-Hochberg correction post-hoc test. $P<0.001$ considered statistically significant.

#### Supplemental Figure 13

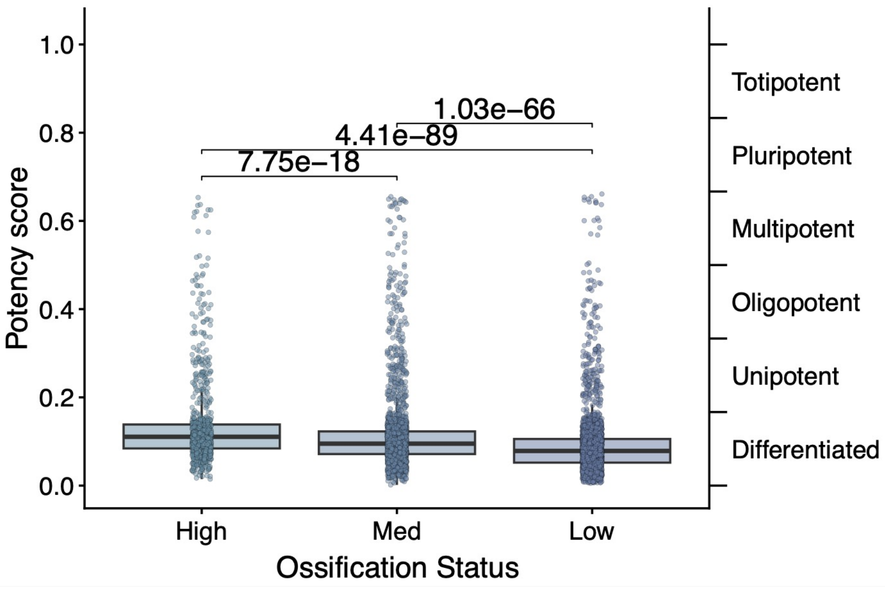

**Supplemental Figure 14: Principal component analysis of bulk RNA-sequencing data from the endothelial co-culture assay.** Principal component analysis (PCA) graphs showing the variability attributed to PC1 vs. PC2 (left) and PC2 vs. PC3 (right). Each dot represents an individual biological replicate and are colored according to the legend to the right of each graph. The graph on the left shows that PC1 explains most of the variation and that this is driven by endothelial cells co-cultured with either OS186 or OS742 osteosarcoma cells. In the right graph, plotting PC2 and PC3 shows that endothelial cells cultured in endothelial growth media are distinct from endothelial cells cultured in other experimental conditions.

**Supplemental Figure 14:**

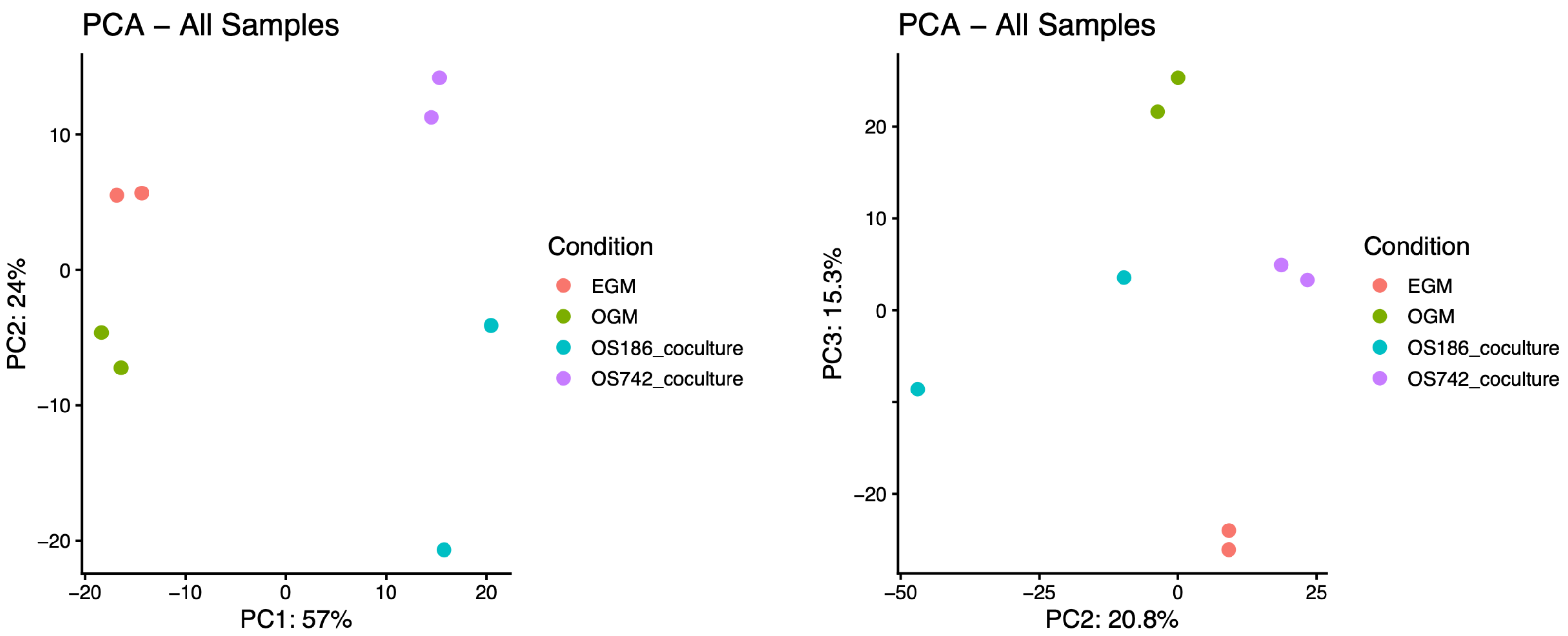
