## Supplemental Table1 for "Integrated multi-omic analysis of pediatric metastatic osteosarcoma reveals endothelial cell plasticity and lineage infidelity"

| Sample_Name | Disease Status | Tissue | Age | Treatment Status |
|---|---|---|---|---|
| CCDI0054n59_1535 | Metastatic | Para-aortic Lymph Node | 12 | post-treatment |
| CCDI0046n64_1600 | Metastatic | Liver | 17 | post-treatment |
| CCDI0094_7977 | Metastatic | Lung | 16 | post-treatment |
| CCDI0095_7978 | Metastatic | Lung | 16 | post-treatment |
| CCDI0090_2318 | Metastatic | Lung | 11 | post-treatment |
| CCDI0048_5864 | Metastatic | Lung | 14 | post-treatment |
| CCDI0049_5899 | Metastatic | Lung | 14 | post-treatment |
| CCDI0050_5928 | Metastatic | Chest Wall | 14 | post-treatment |
| CCDI0051_6547 | Metastatic | Lung | 17 | post-treatment |
| CCDI0091_3666 | Metastatic | Lung | 16 | post-treatment |
| CCDI0092_3911 | Metastatic | Lung | 16 | post-treatment |
| CCDI0093_6546 | Metastatic | Lung | 17 | post-treatment |
| CCDI0099_5913 | Metastatic | Chest Wall | 14 | post-treatment |
| CCDI0100_12416 | Metastatic | Lung | 21 | post-treatment |
| CCDI0096_8952 | Metastatic | Lung | 10 | post-treatment |
| CCDI0098_8951 | Metastatic | Lung | 10 | post-treatment |
| CCDI0097_8978 | Metastatic | Lung | 21 | post-treatment |
| CCDI0052_11487 | Metastatic | Superior Sulcus | No data | No data |
| CCDI0053_445 | Metastatic | Lung | 15 | No data |
| CCDI0055_1804 | Metastatic | Mediastinum | 12 | post-treatment |
| CCDI0057_16178 | Metastatic | Lung | 11 | No data |
| CCDI0058_16183 | Metastatic | Lung | 11 | No data |
| CCDI0101_16178b | Metastatic | Lung | 11 | No data |
| CCDI0102_16183b | Metastatic | Lung | 11 | No data |
