## Supplemental Table 2 for "Integrated multi-omic analysis of pediatric metastatic osteosarcoma reveals endothelial cell plasticity and lineage infidelity"

| Reagent | RRID | Vendor |
| --- | --- | --- |
| 20X Nuclei buffer | --- | 10X Genomics |
| Goat Anti-GFP Polyclonal Antibody | RRID:AB_305643 | Abcam |
| Alvetex 3D scaffolds Amsbio AMS.AVP004-96 | --- | Ambbio |
| TdTomato (E3G5L) Rabbit Monoclonal Antibody | RRID:AB_3662938 | Cell Signaling Technologies |
| 5M NaCl | --- | Corning |
| 3 mM cell insert for 6-well plate | --- | Corning |
| Gelatin coated 6-well plate | --- | Corning |
| Matrigel, Phenol red-free, growth factor reduced | --- | Corning |
| HyClone Bovine Growth Serum (BGS) | --- | Cytiva |
| Bovie Serum Albumin (BSA) | --- | GoldBio |
| 1M $MgCl_2$ | --- | Invitrogen |
| 5% Digitonin | --- | Invitrogen |
| PBS | --- | Invitrogen |
| IMDM, GlutaMAX Supplement | --- | Invitrogen |
| Penicillin-Streptomycin (5,000U/mL) | --- | Invitrogen |
| GelTrex LDEV free reduced growth factor (15 mg/ml) | --- | Invitrogen |
| Opti-MEM Reduced Serum Medium | --- | Invitrogen |
| Hank's Balanced Salt Solution, no calcium, no magnesium | --- | Invitrogen |
| DMEM, high glucose, GlutaMAX Supplement | --- | Invitrogen |
| AO/PI | --- | Logos Biosystems |
| OGM Osteoblast Growth Medium BulletKit | --- | Lonza |
| EGM-2 MV Microvascular Endothelial Cell Growth Medium-2 BulletKit | --- | Lonza |
| Human lung microvascular endothelial cells (HMVEC-L) | --- | Lonza |
| 1M DTT | --- | Millipore Sigma |
| RNase inhibitor | --- | Millipore Sigma |
| 4-hydroxytamoxifen | --- | Millipore Sigma |
| gentleMACS C tube | RRID:SCR_020270 | Miltenyi |
| gentleMACS Octo dissociator | RRID:SCR_020271 | Miltenyi |
| MACS SmartStrainers (70 µm) (Miltenyi #130-098-462) | --- | Miltenyi |
| MACS SmartStrainers (30 µm) (Miltenyi #130-098-458) | --- | Miltenyi |
| NEBNext Ultra II Directional RNA Library Prep Kit for Illumina | --- | New England Biolabs |
| NEBNext Poly(A) mRNA Magnetic Isolation Module | --- | New England Biolabs |
| NEBNext® Multiplex Oligos for Illumina® (96 Unique Dual Index Primer Pairs) | --- | New England Biolabs |
| Fetal Bovine Serum (FBS), Heat Inactivated | --- | Omega Scientific |
| 1M Tris-HCl pH 7.5 | --- | Quality Biological |
| Nuclease-free $H_2O$ | --- | Quality Biological |
| Recombinant Human BMP-4 Protein | --- | R&D Systems |
| Corn Oil | --- | SelleckChem |
| Formical-4 Decalcifier | --- | StatLab |
| OS186 Human Osteosarcoma Cell Line | RRID:CVCL-C8FT | Alejandro Sweet-Cordero (UCSF) |
| OS742 Human Osteosarcoma Cell Line | RRID:CVCL_C8FY | Alejandro Sweet-Cordero (UCSF) |
| OS052 Human Osteosarcoma Cell Line | RRID:CVCL_C8FR | Alejandro Sweet-Cordero (UCSF) |
| OS384 Human Osteosarcoma Cell Line | RRID:CVCL_C8FU | Alejandro Sweet-Cordero (UCSF) |
| BLOXALL Blocking Solution | RRID:AB_2336257 | Vector Laboratories |
| Normal Horse Serum Blocking Solution | RRID:AB_2336617 | Vector Laboratories |
| Antigen Unmasking Solution, Citric Acid Based | RRID:AB_2336226 | Vector Laboratories |
| ImmPRESS-HRP Anti-Rabbit IgG Polymer Detection Kit | RRID:AB_2336529 | Vector Laboratories |
| ImmpACT 3,3'-diaminobenzidine | RRID:AB_2336520 | Vector Laboratories |
| ImmPRESS-AP Anti-Goat IgG (alkaline phosphatase) Polymer Detection Kit | RRID:AB_2828010 | Vector Laboratories |
| ImmPACT Vector Red Substrate Kit | RRID:AB_2336524 | Vector Laboratories |
| Tween-20 | --- | VWR |
| NP40 | --- | VWR |
| 10% neutral buffered formalin | --- | VWR |
| CDH5-$Cre^{ERT2}$; Rosa26-tdTomato | --- | Yosuke Mukoyama (NHLBI) |
| C57BL/6J mice | RRID:IMSR_JAX:000664 | Jackson Laboratories |
| AXT Cells | --- | Takatsune Shimizu (Hoshi University) |
| RNA/DNA/Protein Purification Plus kit | --- | Norgen Biotek |
