## Supplemental Table 3 for "Integrated multi-omic analysis of pediatric metastatic osteosarcoma reveals endothelial cell plasticity and lineage infidelity"

| **Packages** | RRID |
|---|---|
| BSgenome_1.76.0 | RRID:SCR_024230 |
| BSgenome.Hsapiens.UCSC.hg38_1.4.5 | RRID:SCR_027738 |
| cellranger-arc 2.0.0 | RRID:SCR_023897 |
| chromvar_1.30.1 | RRID:SCR_026570 |
| ComplexHeatmap_2.24.1 | RRID:SCR_017270 |
| CytoTRACE2_1.1.0 | RRID:SCR_022828 |
| dittoSeq_1.20.0 | |
| dplyr_1.1.4 | RRID:SCR_016708 |
| DRAGEN BCL Convert 4.2.7 | |
| EnhancedVolcano_1.29.1 | RRID:SCR_018931 |
| EnsDb.Hsapiens.v86_2.99.0 | |
| ensembldb_2.32.0 | RRID:SCR_019103 |
| epiAneuFinder | RRID:SCR_026269 |
| GenePattern | RRID:SCR_003201 |
| ggplot2_4.0.1 | RRID:SCR_014601 |
| ggraph_2.2.2 | RRID:SCR_021239 |
| GraphPad Prism_10.6.1 | RRID:SCR_002798 |
| harmony_1.2.4 | RRID:SCR_022206 |
| igraph_2.2.1 | RRID:SCR_019225 |
| IRanges_2.42.0 | RRID:SCR_006420 |
| JASPAR2020_0.99.10 | RRID:SCR_003030 |
| leidenAlg_1.1.5 | |
| MACS2_2.2.9.1 | RRID:SCR_013291 |
| Morpheus by Broad Institute | RRID:SCR_017386 |
| motifmatchr_1.30.0 | RRID:SCR_026739 |
| msigdbr_25.1.1 | RRID:SCR_022870 |
| paletteer_1.7.0 | |
| patchwork_1.3.2 | RRID:SCR_024826 |
| plyr_1.8.9 | RRID:SCR_026985 |
| presto_1.0. | |
| RColorBrewer_1.1-3 | RRID:SCR_016697 |
| RENEE 2.6.7 | |
| RSEM 1.3.3 | RRID:SCR_000262 |
| Scanpy 1.8.2 | RRID:SCR_018139 |
| scCustomize_3.2.2 | RRID:SCR_024675 |
| SCENIC+ 1.0a2 | RRID:SCR_026702 |
| SCEVAN_1.0.1 | |
| scrublet_0.2.3 | RRID:SCR_018098 |
| scType | RRID:SCR_026634 |
| Seurat_5.3.1 | RRID:SCR_016341 |
| SeuratExtend_1.2.5 | RRID:SCR_026143 |
| Signac_1.16.0 | RRID:SCR_021158 |
| SingleCellExperiment 1.24.0 | RRID:SCR_026794 |
| Snakemake 8.5.5 | RRID:SCR_003475 |
| SoupX_1.6.2 | RRID:SCR_019193 |
| STAR 2.7.6a | RRID:SCR_004463 |
| TFBSTools_1.46.0 | RRID:SCR_024260 |
| tibble_3.3.1 | RRID:SCR_026493 |
| tidygraph_1.3.1 | RRID:SCR_027617 |
| tidyr_1.3.2 | RRID:SCR_017102 |
| tidyverse_2.0.0 | RRID:SCR_019186 |
| UCell_2.12.0 | RRID:SCR_027109 |
| viridis_0.6.5 | RRID:SCR_016696 |
