## Supplemental Table 4 for "Integrated multi-omic analysis of pediatric metastatic osteosarcoma reveals endothelial cell plasticity and lineage infidelity"

| Cell Type | Gene List |
|---|---|
| Osteoclasts | ACP5, CTSK, ITGB3, MMP13, MMP9 |
| Osteosarcoma Cells | RUNX2, ALPL, SP7, SATB2, CDH11, IBSP, COL1A1, SOX9, ACAN, PTH1R, COL27A1, VDR, POSTN, ASPN, VIM |
| Myeloid Cells | SPI1, LGALS1, COTL1, MS4A6A, SRGN, TYROBP, LYZ, FCER1G, LAPTM5, AIF1 |
| T Cells | CD3D, CD3E, CD3G, CD8A, CD2, TRAC, CD4 |
| Smooth Muscle Cells | DES, SOD3, MYL9, ACTA2, MGP, CALD1, TPM1, TAGLN, IGFBP7, TPM2 |
| B Cells | CD79A, MS4A1, CD79B, BANK1, CD37, CD22 |
| Fibroblasts | BGN, DCN, MGP, SPARC, CALD1, LUM, COL6A1, IGFBP7, COL1A2, C1S, ACTA2, FAP, VIM |
| Erythroctye Precursors | HBG2, HBA1, HBA2, HBG1, ALAS2, SPTA1 |
| Lung Epithelial Cells | SFTPB, SFTPC, SFTA3, WIF1, SFTPA1, EPCAM, CDH1 |
| Blood EC | PTRF, CLDN5, AQP1, PECAM1, NPDC1, VWF, RAMP3, RAMP2, SPARCL1, CLEC14A, CDH5, ACKR1 |
| Lymphatic EC | PPFIBP1, GNG11, RAMP2, CCL21, MMRN1, IGFBP7, SDPR, TM4SF1, CLDN5, ECSCR, FLT4, PLVAP |
