## Supplemental Table 5 for "Integrated multi-omic analysis of pediatric metastatic osteosarcoma reveals endothelial cell plasticity and lineage infidelity"

**tdTomato PCR:**

| PCR Cycling Conditions | | | |
|---|---|---|---|
| Step 1: | 94C for 2 min | | |
| Step 2: | 94C for 20 sec | 10 cycles | |
| Step 3: | 65C for 15 sec (-0.5C per cycle) | | annealing |
| Step 4: | 68C for 10 sec | | extention |
| Step 5: | 94C for 15 sec | 28 cycles | |
| Step 6: | 60C for 15 sec | | annealing |
| Step 7: | 68C for 10 sec | | extention |
| Step 8: | 68C for 3 min | | |
| Step 9: | 4C hold | | |

| **Primers** | **Sequence** | **Amplicon Size** |
|---|---|---|
| IMR9020 | AAG GGA GCT GCA GTG GAG TA | 297bp (WT) |
| IMR9021 | CCG AAA ATC TGT GGG AAG TC | |
| IMR9103 | GGC ATT AAA GCA GCG TAT CC | 196 bp (Mutant) |
| IMR9105 | CTG TTC CTG TAC GGC ATG G | |

**CreER PCR:**

| PCR Cycling Conditions | | | |
|---|---|---|---|
| Step 1: | 94C for 2 min | | |
| Step 2: | 94C for 15 sec | 30 cycles | |
| Step 3: | 60C for 15 sec | | annealing |
| Step 4: | 68C for 15 sec | | extention |
| Step 5: | 68C for 3 min | | |
| Step 6: | 4C hold | | |

| **Primers** | **Sequence** | **Amplicon Size** |
|---|---|---|
| Cre FWD | GCGGTCTGGCAGTAAAAACTATC | 100bp |
| Cre RVS | GTGAAACAGCATTGCTGTCACTT | |
