## Supplemental Methods for "Integrated multi-omic analysis of pediatric metastatic osteosarcoma reveals endothelial cell plasticity and lineage infidelity"

### Patient specimens

Archived frozen specimens from pediatric patients with osteosarcoma and clinically confirmed metastatic disease were obtained from the Department of Pathology at Children's Hospital Los Angeles under Institutional Review Board protocol CCI12-00224. Detailed clinical attributes of the specimens are listed in **Supplementary Table S1**.

### Nuclei isolation:

Flash frozen tissue specimens were placed in a mortar in liquid nitrogen and broken into smaller pieces using a pestle. Tissues embedded in OCT were trimmed of excess OCT, placed into phosphate buffered saline (PBS) until the OCT was dissolved, and then the tissue was cut into smaller pieces using a scalpel. The tissues pieces were weighed and placed into a gentleMACS C tube (Miltenyi) containing 1.9ml of NP40 lysis buffer (10mM Tris-HCl pH7.5, 10mM NaCl, 3mM $MgCl_2$, 0.1% NP40, and 1U/µl RNase inhibitor in nuclease-free water). Tissues were homogenized using a gentleMACS Octo dissociator (Miltenyi) using the 4C_nuclei_1 program. Following dissociation, tubes were incubated on ice for 5min. The dissociated tissue was then filtered through a 70µm cell strainer (Miltenyi) into a new tube on ice. Samples were spun for 5min at 500g at 4°C, all but 100ul of the supernatant was transferred to a separate tube, and 900µl of a solution containing 1% bovine serum albumin (BSA) and 0.4U/µl RNase inhibitor (diluted in PBS) was added to the pellet without resuspending. After 5min incubation on ice, the pellet was resuspended in the BSA/RNase inhibitor solution. The samples were then spun again for 5min at 500g at 4°C, and the supernatant was discarded. Pellets were resuspended in 900ul of the BSA/RNase inhibitor/PBS solution and filtered through a 30µm strainer (Miltenyi) into a new tube. Nuclei were counted using acridine orange/propidium iodide dye (AO/PI), and a minimum of 4.5 $x10^5$ nuclei were transferred to a new tube. After spinning the nuclei at 500g at 4°C for 5min, the supernatant was discarded, then 100ul of 0.1X lysis buffer (100ul of 1X lysis buffer (10mM Tris-HCl pH7.5, 10mM NaCl, $MgCl_2$,

0.1% Tween-20, 0.1% NP40, 0.01% digitonin, 1% BSA, 1mM DTT, 1U/µl RNase inhibitor in nuclease-free water) diluted with 900ul of lysis dilution buffer (10mM Tris-HCl pH7.5, 10mM NaCl, 3mM $MgCl_2$, 1% BSA, 1mM DTT, and 1U/µl RNase inhibitor in nuclease-free water) was added to permeabilize the nuclei, and the pellet was gently resuspended. Nuclei suspensions were incubated on ice for 2min, then 950ml of wash buffer (10mM Tris-HCl pH7.5, 10mM NaCl, 3mM $MgCl_2$, 1% BSA, 0.1% Tween-20, 1mM DTT, 1U/µl RNase inhibitor in nuclease-free water) was added and briefly mixed by gentle pipetting. Nuclei were spun at 500g for 5min at 4°C, and the supernatant was discarded. Nuclei pellets were then resuspended in diluted nuclei buffer (1X Nuclei buffer (10x Genomics), 1mM DTT, and 1U/µl RNase inhibitor in nuclease-free water) so that the final concentration of nuclei was at least 5,000 nuclei/µl. Nuclei suspensions were transferred to DNA LoBind microfuge tubes (Eppendorf) and submitted to the POB-CCDI core facility for capture using the 10X Chromium platform.

**Nuclei capture and library prep:**

Upon receipt of the isolated nuclei by the core facility, the quantity and quality of the single nuclei suspension was assessed using an AO/PI stain. Typically, samples with more than 5% viability were excluded. However, to avoid decreasing the sample size of this study, samples that included small fragments of osteoid were included. As these samples had higher false positive background, the loading concentrations were calculated using the number of dead cells per the AO/PI results, adjusting to 4,000-5,000 nuclei/µl. The quality and the concentrations were again checked and then the single nuclei capture with a targeted recovery of 7,000 nuclei, was performed by following the 10X Genomics User Guide (CG000338). Briefly, the open chromatin regions of the DNA were transposed and then the adapter sequences were added to capture the DNA fragments for ATAC library. The transposed nuclei along with RT mix and beads were added on to the NextGEM Chip J and loaded immediately to the Chromium Controller for GEM generation and barcoding. After the Post-GEM cleanup and pre-amplification PCR, 10X barcoded DNA fragments and the 10X barcoded full length cDNA from mRNA in the pre-amplified product were used to prepare ATAC library and GEX library respectively. The libraries were prepared as dictated by the 10X Genomics User Guide.

**Sequencing:**

The concentration and average size of the ATAC and GEX final libraries were quality checked using the Agilent 2100 Bioanalyzer (Agilent). Equimolar concentrations of the libraries from all the samples were pooled and sequenced using the NovaSeq-6000 (Illumina). The ATAC and GEX libraries were pooled separately and sequenced at the sequencing depth of 50,000 reads per nucleus. The sequencing parameters are as follows: GEX Library - 28|10|10|90, R1|I1|I2|R2 and ATAC library - 50|8|24|49, R1|I1|I2|R2)

**Data Processing:**

Raw BCL files were first demultiplexed and then aligned to the human reference genome provided by 10X genomics (refdata-cellranger-arc-GRCh38-2020-A-2.0.0). The demultiplexing and alignment were performed using the cellranger-arc pipeline, version 2.0.0. The count matrix files generated for each sample were used for downstream data analyses. The raw gene expression and chromatin accessibility data was loaded into Seurat (v5.0.2) and Signac (1.12.9004) for additional pre-processing, including filtering, normalization, dimensionality reduction, batch correction, clustering. To mitigate the effects of ambient RNA contamination, raw counts of the RNA assay were corrected using SoupX **,** with a fixed contamination rate of 0.20, producing an adjusted gene expression matrix. The corrected count matrix was then used to create a Seurat object for downstream analysis. All QC metrics were computed on a per-sample basis prior to integration. The following criteria were applied to each sample to remove low-quality cells. Only nuclei passing both RNA and ATAC QC thresholds were retained, ensuring high-quality paired multiomic profiles. After filtering, we obtained a dataset of 134,669 nuclei from a total of 24 samples.

*<u>scRNA-seq quality control</u>*

For the RNA modality, nuclei were required to meet the following criteria:

- Total RNA counts (nCount_RNA): ≥ 500 and ≤ an upper threshold defined per sample using a median absolute deviation (MAD)–based outlier detection (3 MADs above the median).
- Number of detected genes (nFeature_RNA): > 300.
- Mitochondrial gene fraction: < 10% of total UMI counts.
- Doublets were identified using scrublet, a simulation-based method that assigns each cell a doublet score based on synthetic doublet profiles. For each sample, a dynamic doublet score threshold was calculated based on the expected doublet rate 4%, scaled by the number of recovered cells. This adaptive approach ensured consistent doublet filtering across samples with varying cell recoveries.

*scATAC-seq quality control*

For the ATAC modality, nuclei were required to meet the following criteria:

- Nucleosome signal: < 2.
- TSS enrichment score: > 2.
- Fraction of fragments overlapping ENCODE blacklist regions: < 5%

**Downstream processing of RNA assay data:**

After filtering, gene expression counts log normalized with a scale factor of 10,000. Highly variable genes were identified, and the top 5,000 variable features were retained for downstream analysis. Gene expression values were then scaled across all genes. Cell cycle phase scores were computed using canonical S and G2/M gene sets. Cell cycle scores, along with additional technical covariates including mitochondrial read percentage, ribosomal read percentage, total UMI counts, number of detected genes, and doublet scores, were regressed out during downstream normalization steps. The top 2,000 genes that show the most consistent biological variation across all dataset were selected as integration features using SelectIntegrationFeatures function in Seurat. These features were used for scaling and principal component analysis (PCA) within each sample. To correct for batch effects and enable joint analysis across samples, the RNA assay was integrated using Seurat's reciprocal PCA (RPCA)–based integration workflow. Integration anchors were identified using the selected 2000 variable features, with anchor identification performed across 50 principal components and k.anchor set to 7. The integrated expression matrix was constructed using these anchors and a weighting parameter of k.weight = 50.

**Downstream processing of ATAC assay data:**

ATAC peaks were re-called independently for each sample using MACS2 on the ATAC fragments. Peaks located on nonstandard chromosomes or overlapping ENCODE hg38 blacklist regions were removed. A unified consensus peak set was generated by merging peaks across all samples and reducing overlapping intervals. Consensus peaks were further filtered to retain peaks between 20 bp and 10 kb in width. This consensus peak set was used to quantify chromatin accessibility uniformly across all cells. MACS2-derived chromatin accessibility counts were used for further analysis. Term frequency–inverse document frequency (TF–IDF) normalization was applied to the peak-by-cell matrix to account for sequencing depth and sparsity. Each sample was then processed independently to identify highly informative peaks using FindTopFeatures and top 95% most common features was used in the analysis. A unified feature set was constructed by taking the union of variable peaks across all samples. Linear dimensionality reduction was performed using latent semantic indexing (LSI) and Singular value decomposition (SVD) was applied to the TF–IDF–normalized matrix using the selected variable peaks to generate LSI embeddings. To correct for batch effects and enable joint analysis across samples, the ATAC data was integrated using a reciprocal LSI (RLSI)–based integration workflow. The first LSI component—primarily associated with sequencing depth—was excluded, and subsequent 49 components were used for integration. Integration anchors were identified across samples using the shared variable peak set, with a k.anchor value of 20 and a filtering parameter of k.filter = 100. Integrated embeddings were computed using weighted anchor correction (k.weight = 50) and stored as a unified integrated LSI reduction.

**Joint integration of scRNA-seq and scATAC-seq data**

Joint analysis of single-cell gene expression and chromatin accessibility data was performed using Seurat's weighted nearest neighbor (WNN) framework. For the gene expression modality, the integrated RNA assay was used as input. Gene expression values were scaled, and principal component analysis (PCA) was performed using all genes. A nearest neighbor graph was constructed based on 50 principal components. For the chromatin accessibility modality, previously computed integrated latent semantic indexing (LSI) embeddings were used, excluding the first LSI component to mitigate sequencing depth–associated effects. Weighted nearest neighbor (WNN) analysis was performed using PCA embeddings from the RNA modality and integrated LSI embeddings from the ATAC modality to calculate multimodal neighbor graphs with the number of neighbors used to compute modality weights k.nn set to 100. Clustering was performed on the weighted shared nearest neighbor (wsnn) graph using the SLM algorithm across a range of resolutions (0.1-1). A final clustering resolution of 0.1 was selected for downstream analyses, and clusters with cell number <= 100 were excluded in the further analysis. A joint low-dimensional embedding was generated using UMAP based on the weighted nearest neighbor graph.

**Copy Number Alterations (CNAs) analysis from scRNA-seq and ATAC-seq data**

CNA analysis was performed on the scRNA-seq data using SCEVAN, which estimates large-scale chromosomal copy number alterations by leveraging relative gene expression patterns along genomic coordinates at single cell resolution. Based on marker genes for myeloid cells, T cells, and lung epithelial cells, we defined a set of confident normal cells which are used as normal reference in SCEVAN. For each sample, raw RNA count matrices were used as input for SCEVAN to classifies cells into malignant and non-malignant types.

For scATAC-seq data, CNAs were inferred using epiAneufinder by quantifying read counts in genomic bins and identifying segmented gains and losses across single cells, enabling the detection of large-scale copy number alterations from scATAC-seq data alone. For each sample, scATAC-seq fragment files generated by Cell Ranger ARC were used as input. Cells with fewer than 4000 ATAC fragments were excluded from CNA analysis to reduce sparsity and technical noise. Fragment counts were aggregated into fixed 2Mb genomic bins. Only autosomal chromosomes were included, while sex chromosomes (chrX, chrY) and mitochondrial DNA (chrM) were excluded from analysis. Genomic regions overlapping ENCODE blacklist regions were also excluded from the analysis.

**Subsetting and processing of endothelial cell nuclei and vascular mimics**

The annotated endothelial cell nuclei were subsetted from the larger dataset. ATAC peak calling was re-run on the endothelial subset to capture cell-type specific chromatin accessibility that would otherwise be masked in the full dataset. Subsequently, processing and integration of the RNA and ATAC data was performed as described above. The nuclei were clustered using the Louvain algorithm in Seurat and cluster-specific marker genes were identified.

The endothelial dataset was further subset based on the CNV status. Nuclei labeled normal by both SCEVAN and epiAneuFinder were annotated diploid endothelial cell nuclei while nuclei labeled as tumor by both tools were annotated as osteosarcoma vascular mimics. Nuclei that were labeled tumor by onr tool and normal by the other were labeled as ambiguous. Batch correction was performed on the RNA assay and the MACS assay using Harmony prior to WNN integration. The individual assays were clustered using Louvain algorithm.

**Gene regulatory network (GRNs) analysis in endothelial cells**

GRNs were built using SCENIC+ (v1.0a2), following the tutorial at https://scenicplus.readthedocs.io/en/latest/index.html. Briefly, gene expression matrix and chromatin accessibility counts were converted into h5ad format separately using SingleCellExperiment (v1.24.0), which was recognizeable by Scanpy (v 1.8.2). To generate a custom cisTarget database, which is recommended by the tutorial, consensus ATAC-seq peaks with extension of ±1 kb was used to extract sequences from the GRCh38 reference genome. The RNA data and ATAC fragments data were used to create the cisTopic object. Chromatin accessibility Topic modeling for scATAC-seq data was performed using pycisTopic with the MALLET implementation and 25 Topics was selected for downstream analysis. Subsequently, the config.yaml file was completed with relevant modification and the SCENIC+ workflow was ran using Snakemake (v8.5.5) in multiome mode. The AUCell matrix for eRegulons from the output was added into the endothelial cell Seurat object as assays for further visualization and downstream analysis.

**Bulk RNA sequencing**

Poly- A libraries were constructed from 800 ng of total RNA from each sample using the NEBNext Ultra II Directional RNA Library Prep kit for Illumina (NEB) as per the manufacturer's protocol. Sequencing was performed on the NextSeq2000 (Illumina) with a 101x101 pair end configuration. The demultiplexing was performed using the DRAGEN BCL Convert (v 4.2.7) workflow. The demuliplexed data were processed using the RENEE pipeline (RNA-sEq aNalysis pipeline, v2.6.7) available through the CCR Collaborative Bioinformatics Resource (CCBR) at the National Cancer Institute. Raw paired-end FASTQ files were aligned to the human reference genome hg38 using the hg38_45 reference bundle (GENCODE release 45) with STAR v2.7.6a. Gene-level quantification was performed using RSEM (v1.3.3) to generate gene-level expected counts.

**Chromvar**

Chromvar (v1.30.1) was used to analyze transcription factor motif activity by following the vignette on the Seurat website (https://satijalab.org/seurat/articles/weighted_nearest_neighbor_analysis). First, the default assay was set to "RNA" and the layers of the object were joined using "JoinLayers", after which the default assay was set to "MACS". The GC content was computed for each peak with "RegionStats" then the DNA sequence was scanned to find transcription factor binding motifs present according to the JASPAR 2020 database. The motif matrix and subsequent motif objects were created using the “CreateMotifMatrix” and “CreateMotifObject” functions. Motif activity UMAP plots were generated by first converting the gene name into a motif ID using “ConvertMotifID”, then plotted using the “FeaturePlot_scCustom” function in the scCustomize package.

**Gene set enrichment analysis**

The UCell package (v2.12.0) was used to perform gene set enrichment on gene signatures from the Reactome, Gene Ontology: Biological Processes, and Wikipathways libraries within the Molecular Signatures Database (MSigDB). The metagene signature scores were added to the Seurat object metadata using the “AddModuleScore_UCell” function. Lung endothelial cell (Lung-EC) and bone marrow mesenchymal cell (BM-MC) signatures were derived from the Tabula Sapiens dataset. Marker genes from normal bone marrow mesenchymal cells and normal lung endothelial cells with the highest effect size and specificity score were selected and queried against the subsetted endothelial cell nuclei from the snMulti-ome dataset to ensure their expression. Marker genes not expressed in the snMulti-ome dataset were removed from the gene list.

**Cell type annotations**

ScType was used to initially annotate the data using the marker genes in **Supplemental Table 4**. Nuclei annotated as “Unknown” were subclustered using the “FindSubCluster” function and marker genes were identified for each sublcuster using the “FindMarkers” function in Seurat. The top 250 marker genes for each subcluster were submitted to the Enrichr online database and queried against multiple cell type databases including CellMarker 2024, Descartes Cell Types and Tissue 2021, Tabula Sapiens, PanglaoDB Augmented 2021, Tabula Muris, and Azimuth 2023 to determine the cell type. Subclusters were manually re-annotated using the corresponding cellular type when consensus was reached in two or more databases. The cellular annotations produced by scType were further refined by subsetting each annotated cluster and subclustering to ensure that each subcluster expressed marker genes associated with the appropriate cellular lineage. As stated above, the top 250 marker genes for each subcluster were submitted to the Enrichr online database and each subcluster was manually re-annotated according to the resultant consensus across multiple cell type databases. Clusters with small numbers of non-informative nuclei were not annotated and removed from the dataset.

**CytoTRACE2**

Evaluation of stemness/potency was performed using CytoTRACE2 as per the vignette.

**Pseudotime trajectory analysis**

Pseudotime trajectory analysis was performed using the "RunSlingshot" function in the SeuratExtend package as per the vignette.

**Mass spectometry**

*Digestion and TMTpro labeling:* Samples in TPER buffer were precipitated with 100% TCA (4:1 v/v ratio sample:TCA) and resuspended in 500µL Easypep lysis buffer (Thermo #A45735) with 1x phosphatase inhibitors (Thermo #A32957) and briefly sonicated. 120µg was taken from each sample for digestion. Samples were adjusted to 230µL total with lysis buffer and treated with 70µL of digestion buffer (50mM TCEP, 120mM chloroacetamide, and 50ng/µL trypsin/LysC in 100mM HEPES pH 8) and incubated at 37°C in the dark. A reference pool was made using 10µg of each digested sample. Excess TMTpro was quenched with 50µL of 5% hydroxylamine, 20% Formic acid for 30min and samples from each batch were combined along with equal amount of reference pool. Batches were cleaned using EasyPep Maxi columns (Thermo #A45734) and eluted with 3mL of the provided elution buffer. Eluted peptides were aliquoted and dried in speed-vac. For each batch, the eluted peptides from each enrichment were combined and dried.

*LC/MS analysis:* For each batch the protein aliquot was resuspended in 50µL of 0.1%FA and 5µL was analyzed in triplicate. All samples were analyzed using a Dionex U3000 RSLC in front of a Orbitrap Lumos (Thermo) equipped with an EasySpray ion source. Solvent A consisted of 0.1%FA in water and Solvent B consisted of 0.1%FA in 80%ACN. Loading pump consisted of Solvent A and was operated at 7 µL/min for the first 6 minutes of the run then dropped to 2 µL/min when the valve was switched to bring the trap column (Acclaim™ PepMap™ 100 C18 HPLC Column, 3µm, 75µm I.D., 2cm, PN 164535) in-line with the analytical column EasySpray C18 HPLC Column, 2µm, 75µm I.D., 25cm, PN ES902). The gradient pump was operated at a flow rate of 300nL/min and each run used a linear LC gradient of 5-7%B for 1min, 7-30%B for 133min, 30-50%B for 35min, 50-95%B for 4min, holding at 95%B for 7min, then re-equilibration of analytical column at 5%B for 17min. All injections employed the TopSpeed method with three FAIMS compensation voltages (CVs) and a 1 second cycle time for each CV (3 second cycle time total). Global parameters for all methods were the same with the spray voltage set to 2200V and ion transfer temperature of 300 ºC. MS1 scans were acquired in the Orbitrap with resolution of 120,000, AGC of 4e5 ions, and max injection time of 50ms, mass range of 350-1600 m/z; MS2 scans were acquired in the Orbitrap using TurboTMT method with resolution of 15,000, AGC of 1.25e5, max injection time of 22ms, HCD energy of 38%, isolation width of 0.4Da, intensity threshold of 2.5e4 and charges 2-6 for MS2 selection. Advanced Peak Determination, Monoisotopic Precursor selection (MIPS), and EASY-IC for internal calibration were enabled and dynamic exclusion was set to a count of 1 for 15sec. The only difference in the methods was the CVs used. Global peptide analysis used three methods consisting of CVs -45, -60, -75 for method one; -50, -65, -80 for method two; and -55, -70, -85 for method three.

*Database search and post-processing analysis:* All raw files were batched together as fractions and searched with Proteome Discoverer 2.4 using the Sequest node. Data was searched against the Uniprot Human database from Feb 2020 using a full tryptic digest, 2 max missed cleavages, minimum peptide length of 6 amino acids and maximum peptide length of 40 amino acids, an MS1 mass tolerance of 10 ppm and MS2 mass tolerance of 0.02 Da. Variable modifications of oxidation on methionine (+15.995 Da) and phosphorylation on serine, threonine, and tyrosine (+79.966); fixed modifications of carbamidomethyl on cysteine (+57.021) and TMTpro (+304.207) on lysine and peptide N-terminus. Percolator was used for FDR analysis and TMTpro reporter ions were quantified using the Reporter Ion Quantifier node. Proteins were filtered using an FDR cutoff of 1%. Proteins with quantitative values in the reference pool channel of both batches were included in the final list.

*Data normalization and statistical analysis:* To account for TMT batch effects, protein and peptide intensities were normalized using the Internal Reference Standard (IRS) method as described in Plubell et. al. (https://pubmed.ncbi.nlm.nih.gov/28325852/). Briefly, for each TMTpro channel the summed raw intensity of the channels was obtained (sum of all raw intensities in that channel) and then normalized by dividing the highest total intensity of the entire dataset by the total intensity of each channel to obtain the Total Channel Normalized Intensity (TCNI) correction factor and all protein/peptide intensities in a given channel were multiplied by the TCNI correction factor to give the TCNI for each protein/peptide. Next, the average of the TCNI for each protein/peptide in the reference pools in each batch was divided by the TCNI of the individual reference pools to generate the IRS correction factor. Proteins/peptides in each batch were then multiplied by the IRS correction factor for the corresponding batch to give the IRS Normalized Intensity (IRS-NI), which was used as the input for single sample gene set enrichment analysis.
